# Hidden symmetries in network connectivity support ring attractor dynamics in the fly’s neural compass

**DOI:** 10.64898/2026.05.18.725766

**Authors:** Brad K. Hulse, P.B. Aneesh, Sandro Romani, Vivek Jayaraman, Ann M. Hermundstad

## Abstract

Theoretical models can explain how network structure shapes neural computation, but they typically assume idealized connectivity that is inconsistent with the heterogeneous wiring of biological circuits. We address this issue in the Drosophila head-direction system, a recurrent network with ring-attractor dynamics that enable angular velocity integration. The network’s symmetric wiring motifs are reminiscent of classical models, but with additional heterogeneity that should, in principle, destabilize attractor dynamics. Inspired by novel architectures discovered through machine-learning-based optimization, we develop an algorithm that transforms attractor models with symmetric connectivity into functionally equivalent models with heterogeneous connectivity. By replacing each unit with multiple clones that preserve its output, the algorithm embeds hidden symmetries in heterogeneous connectivity, maintaining ring-attractor dynamics and accurate integration. Analysis of multiple fly connectomes provides evidence for duplicated units whose connectivity reflects hidden symmetries, consistent with our theory. Our framework helps reconcile idealized models of neural computation with heterogeneous biological circuits.

## INTRODUCTION

Many essential neural computations rely on the brain’s ability to generate stable population activity patterns that encode latent variables such as spatial position [1], orientation [2], goal location [3], or internal state [4]. Foundational work in theoretical neuroscience can explain how recurrent neural circuits might generate these dynamics ([5–21]; reviewed in [22, 23]). Although it has been possible to test functional predictions of circuit models with neuron-type-specific perturbations and recordings [24–31], the recent generation of multiple large-scale, dense connectomes [32–37] provides an opportunity to reexamine the structural assumptions made by these theories [38, 39]. These assumptions are particularly crucial for models of circuit dynamics that underlie the internal representation of continuous variables such as an animal’s position [17] or orientation [40–42] in its environment, which rely on fine-tuned symmetries in synaptic weights to achieve accurate internal representations.

We examine these theoretical assumptions in the context of the *Drosophila* head direction system [24, 43, 44], a recurrent network whose recent connectomes [38, 39] enable direct comparisons with a special class of theoretical models known as continuous attractor networks ([25, 40, 42, 43, 45–48], reviewed in [22, 23, 49]). These models can maintain persistent activity that tracks an animal’s head direction, but they rely on precise and symmetric connectivity to do so; even small deviations from this idealized connectivity can substantially degrade coding accuracy [40–42, 50–52]. This theoretical requirement for strict symmetry suggests that continuous attractor networks are exceedingly rare in the space of all possible networks and may therefore be biologically implausible to implement. In contrast, biological attractor networks are remarkably robust, maintaining accurate function despite substantial variability in wiring [39, 40]. How can the robustness of biological attractors be reconciled with the fine-tuned synaptic connectivity required by canonical attractor models?

Here we use machine-learning-based optimization to explore the space of attractor architectures. In doing so, we discover architectures with heterogeneous connectivity that nevertheless track head direction. Inspired by these unorthodox solutions, we develop a construction algorithm that replaces each unit in canonical attractor networks with multiple “dynamical clones” that preserve the original unit’s output weights. We show how these groups of dynamical clones permit variability in network wiring without altering function, allowing us to transform continuous attractor networks with symmetric connectivity into an infinite set of functionally equivalent networks with heterogeneous connectivity. This algorithm makes clear how symmetric connectivity can be hidden within heterogeneity, and how these hidden symmetries can be recovered. Guided by this understanding, we analyze multiple fly connectomes and find evidence for hidden symmetries in the recurrent connectivity of neurons with shared directional tuning. Together, our results relax the fine-tuning constraints in continuous attractors and help reconcile their connectivity requirements with seemingly messy biological connectomes.

## RESULTS

To accurately encode a continuous angular variable like head direction, neural activity patterns must stably span a continuum of angles and must change in lockstep with the variable being encoded. Classical ring attractor models achieve this via local excitation and long-range inhibition between directionally-tuned neurons (Fig. 1a; [5, 6, 10, 13, 16, 32]). When neurons are ordered on a ring according to their directional tuning, the resulting network connectivity is circularly-symmetric, and the weights can be chosen [40] to support a bump of activity that stably persists at any orientation (Fig. 1b). To move this bump, classical models introduce additional “side” rings of neurons that receive angular velocity input and project back to the “center” ring with phase-shifted connectivity. The resulting connectivity matrix is composed of circularly-symmetric submatrices that define connections within and between rings, with non-zero phase shifts between rings that are characterized by off-diagonal shifts in the connectivity matrix (Fig. 1c) [8, 10]. When appropriately tuned, these side rings support accurate velocity integration (Fig. 1d). Using these key ingredients – rings of neurons with phase-shifted connectivity and velocity modulation – theorists have since proposed several distinct ring attractor architectures with different numbers of rings and velocity integration mechanisms, suggesting multiple potential architectures that biological networks could implement [8–12].

**Figure 1:**
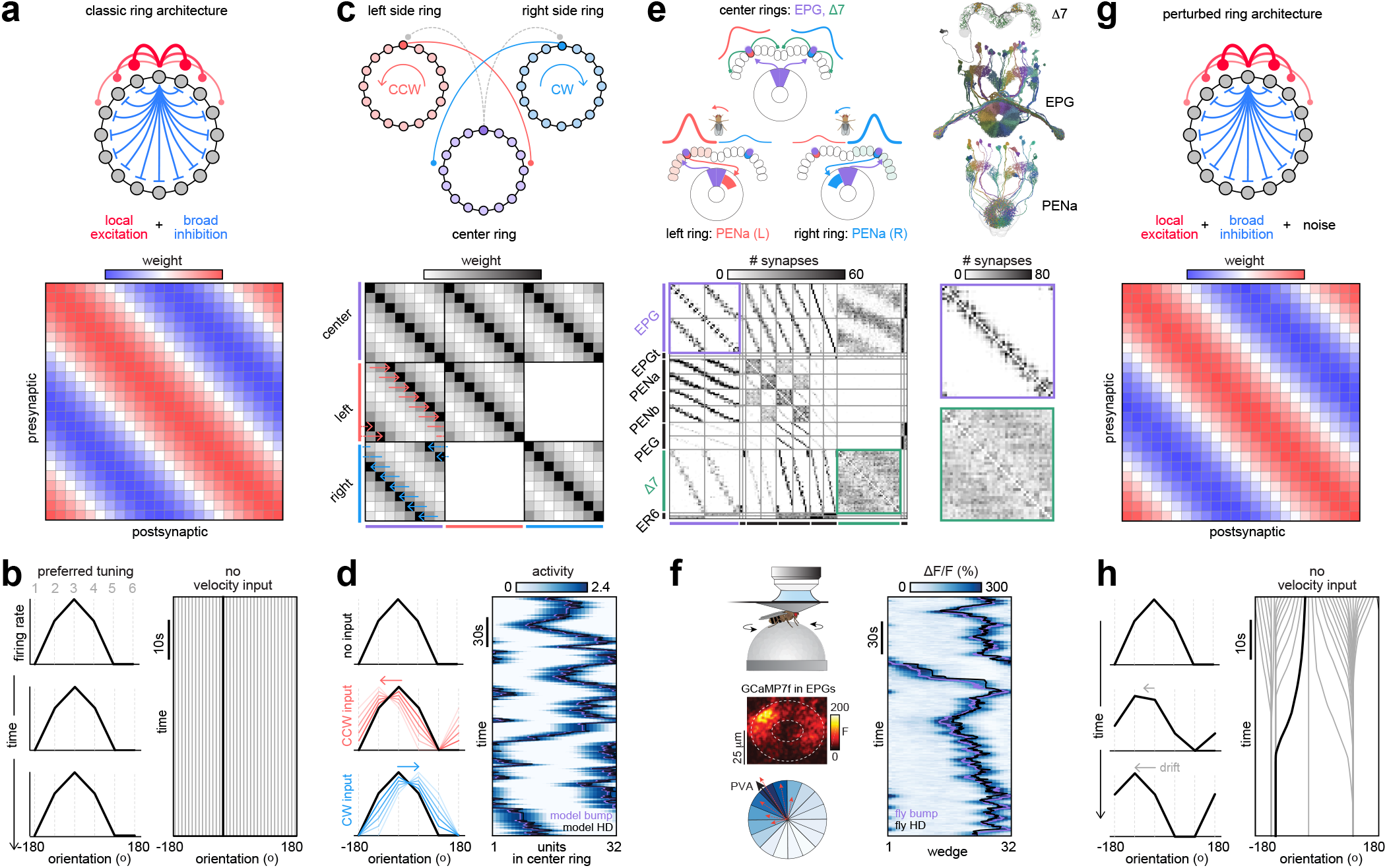
Motivation. **a)** Upper: A continuous ring attractor can be constructed via local excitation (red) and broad inhibition (blue) between neurons that are arranged on a ring according to their tuning to head direction. Lower: The full connectivity matrix is constructed by applying the same profile of excitation and inhibition to all units in the network, so that each successive row is a shifted version of the previous one, making the rows circularly symmetric. **b)** The network shown in panel **a** generates a stable bump of population activity (left column) that can persist at any orientation in the absence of velocity input (right column). **c)** Upper: To move the bump of activity, two side rings inherit the bump from the center ring (gray dashed projections), and project back to the center ring with phase-shifted connections (red and blue projections). When differentially tuned to CW and CCW velocity, the side rings can shift the bump of activity in the center ring. Lower: connections within individual rings are represented by circularly-symmetric submatrices along the diagonal of the full connectivity matrix. Connections between rings are represented by off-diagonal submatrices, which are themselves circularly symmetric but can additionally include phase-shifts (horizontal arrows). **d)** When the connectivity of the network in panel **c** is appropriately tuned, this architecture can accurately integrate angular velocity (right column) by appropriately shifting the bump of activity (left column). **e)** Upper: the fly central complex (CX) contains the core connectivity motifs predicted by theory and shown in panel **c**. Two cell types (EPGs and Δ7s) maintain a bump of activity that tracks the fly’s head direction, while a third cell type (left (L) and right (R) PENa neurons) shifts the bump based on angular velocity input. Lower: Connectivity within EPGs (purple box) and Δ7s (green box) resembles the connectivity of classical ring attractors, but with heterogeneity in synaptic connectivity (compare to the weight matrix in panel **a**). In addition to these motifs, the CX contains many other cell types that could shape attractor dynamics. **f)** Left column: Two photon imaging setup (upper) for tracking EPG activity (middle) in tethered walking flies. We use 32 wedges to compute the population vector average (PVA) of the change in fluorescence (ΔF/F; lower). Right column: The population vector average (purple line) tracks the fly’s head direction (black line) in darkness. **g-h)** Infinitesimal noise in the strength of connections (panel **g**) destroys the continuous attractor (panel **h**; [40, 41]), such that the bump of population activity cannot persist at the same orientation in the absence of velocity input (**h**, right column). This causes the bump to drift over time and settle at a discrete number of stable bump orientations (**h**, left column).

These core connectivity motifs have since found biological support in the fly head direction system (Fig. 1e) [25, 38, 39, 43, 44]. In particular, two neural populations – named EPG and Δ7 – maintain activity bumps that track the fly’s heading and have recurrent connections that are approximately circularly symmetric (Fig. 1e), consistent with theoretical predictions for center rings [24, 38, 39]. Two additional populations – named PENa and PENb – inherit these activity bumps, modulate their amplitudes via angular velocity input [38, 43, 44, 53], and project back to the EPGs with clockwise or counterclockwise phase shifts (Fig. 1e, top schematic; [38, 39, 54]), as predicted by theory [8]. These connectivity motifs ensure that each neural population maintains a localized bump of activity that rotates when the fly turns and persists when the fly stands still, even in darkness (Fig. 1f; [24]). Despite these similarities, the fly circuit is more complex than classical models require, with multiple cell types and potential sources of feedback inhibition that could shape attractor dynamics [39]. In addition – and central to our focus here – the connectivity within and between cells types is heterogeneous, breaking the strict symmetry that is required for theoretical ring attractor networks to maintain and update continuous representations of head direction (compare the heterogeneous Δ7 and EPG connectivity matrices in the lower right panels of Fig. 1e to the perfectly symmetric matrix in Fig. 1a). Indeed, even imperceptible amounts of noise in the synaptic weights (Fig. 1g) can quickly degrade the performance of classical attractor models (Fig. 1h; [40, 41, 55]). This raises the question of how the fly’s head direction circuit can maintain accurate performance despite heterogeneous connectivity.

To address this question, we combine machine learning, theory, and connectomic analyses to explore the space of ring attractor architectures. Using machine-learning-based optimization, we train recurrent neural networks (RNNs) to integrate angular velocity, both with and without connectomic constraints (Fig. 2). Guided by theory, we reverse engineer these RNNs to discover a richer landscape of candidate attractor architectures than classical theory proposes — ones in which units with shared directional tuning differ substantially in their phase-shifted connectivity (Fig. 3). This observation inspired us to develop a construction algorithm that transforms idealized ring attractors with symmetric connectivity into functionally equivalent networks with “hidden” symmetries embedded in heterogeneous connectivity, without sacrificing theoretical guarantees for exact ring attractor dynamics (Figs. 4-5). Understanding how hidden symmetries are constructed allows us to uncover evidence for such symmetries in the recurrent connectivity of Δ7 and EPG neurons across multiple fly connectomes (Fig. 6).

**Figure 2:**
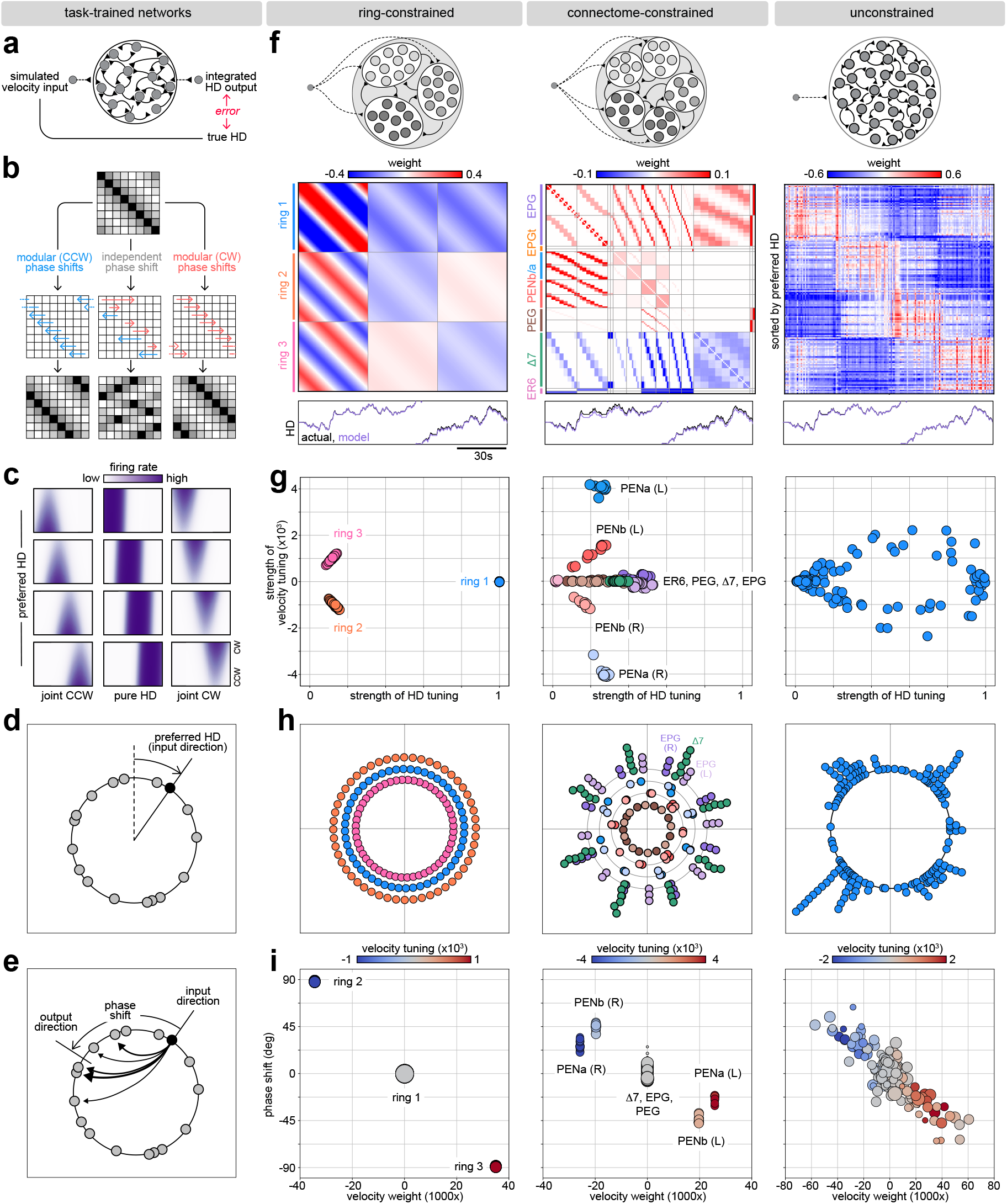
Heterogeneous wiring in trained RNNs. **a)** We train RNNs to integrate angular velocity by minimizing the error between the network output and the true head direction (computed by integrating the network input). **b)** Classical models impose modular constraints on network properties like phase shifts; RNNs can relax these constraints and may learn independent phase shifts for individual units in the network. **c)** Units show a mixture of tuning to HD and angular velocity [58, 59], as observed in HD systems across species [43, 44, 64–67]. Each subpanel shows a single-unit tuning curve to HD (x-axis) and angular velocity (y-axis). For each unit in every network, we measure the HD at which it fires most strongly (“preferred HD”) and the firing rate at that location (“strength of HD tuning”); we also measure the slope of its angular velocity tuning (“strength of velocity tuning”). **d)** We organize units according to their preferred HD (“input direction”). **e)** After organizing units on a ring, we compute the average direction to which each neuron projects (“output direction”). We use the difference between the unit’s input and output direction as a measure of its phase shift. **f)** We impose different classes of structural constraints on trained RNNs (upper row: schematic of RNN; middle row: example connectivity matrix after training; lower row: trained model output for the same velocity input). Ring constrained networks (left column) are constrained to have distinct rings whose connectivity is circularly symmetric; connectome-constrained networks are constrained to preserve the connectivity of the fly compass network (middle column); unconstrained networks do not have any structural constraints (right column). **g)** Different network classes develop different distributions of tuning to HD and angular velocity. Units in ring-constrained (left) and connectome-constrained (middle) networks cluster in the strength of their tuning to HD and angular velocity; units in unconstrained networks (right) span a continuum. **h)** Units in ring-constrained networks (left) evenly tile HD (the three rings are separated radially); units in connectome-constrained (middle) and unconstrained (right) networks cluster in their preferred HD (units with similar tuning have been stacked along the radial direction, with additional grouping by cell type). **i)** Units in ring-constrained (left) and connectome-constrained (middle) networks have clustered phase shifts and velocity weights; units in unconstrained networks (right) span a continuum. See Fig. 2 S1 to S4 for similar plots for the ensemble of 10 networks of each class.

**Figure 3:**
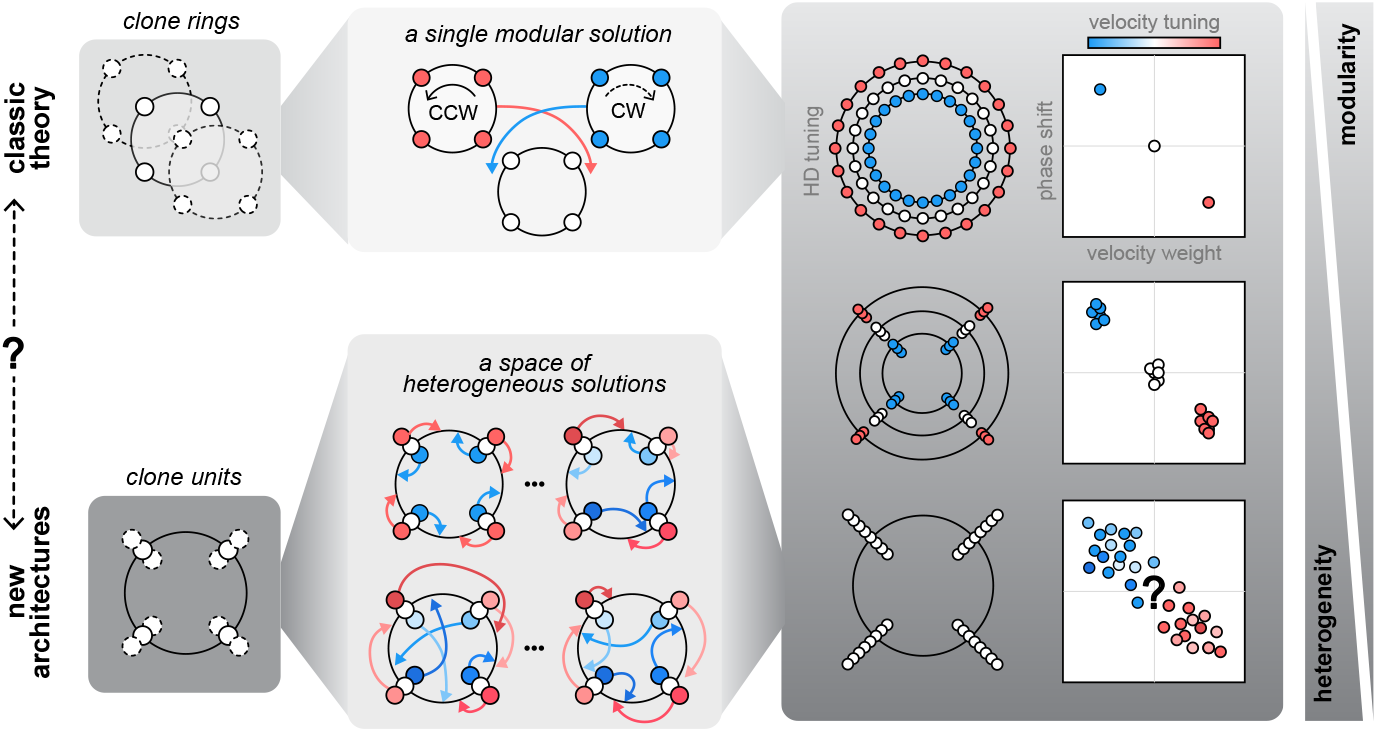
Trained RNNs reveal a space of architectures that vary in their modularity and heterogeneity. Upper row: classical theory typically constructs path-integrating attractor networks by cloning modular rings of neurons [8, 10], and assigning structural and functional properties to individual rings. Each ring preserves a uniform tiling over head directions, and connections between rings share the same phase shifts and velocity weights. Lower row: the solutions discovered by unconstrained RNNs suggest that cloning individual neurons, rather than entire rings, could permit substantially more flexibility in phase shifts and velocity weights.

**Figure 4:**
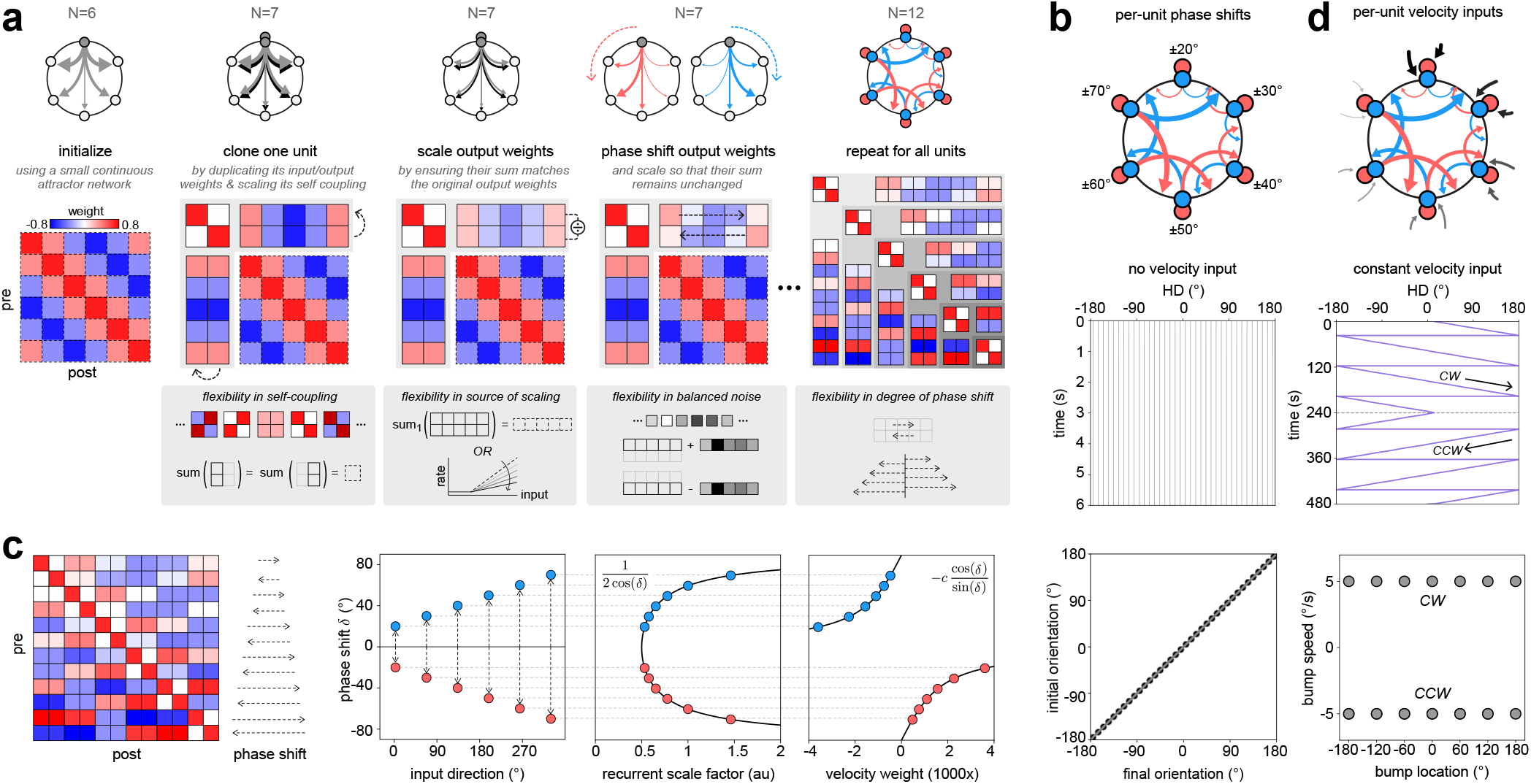
A construction algorithm uses dynamical clones to introduce hidden symmetries in network connectivity. **a)** To construct heterogeneous attractor networks, we begin with a small continuous attractor network whose weights are known (first column; [40]). For each unit in this network, we clone its dynamics; this requires duplicating its input and output weights, and scaling its self-coupling (second column). We then scale the output weights in proportion to the number of cloned units; this can be achieved by weakening the synaptic weights or reducing the slope of the activation function (third column). Once cloned, these units can support a continuum of phase shifts and additive noise that are counterbalanced across units (fourth column). This cloning procedure can be repeated any number of times and across any subset of neurons in the original network to generate an equivalent attractor network with heterogeneous weights and phase shifts (fifth column, shown for 2 clones per unit). **b)** The cloned network maintains a bump of activity that can persist at any orientation in the absence of input. **c)** To accurately integrate angular velocity, recurrent and input weights must be scaled in proportion to the per-unit phase shifts. For the network constructed in panel (a), these phase shifts increase in magnitude across pairs of units (first and second columns). These phase shifts determine the recurrent and velocity input weights (third and fourth columns, respectively). **d)** When recurrent and velocity input weights are appropriately scaled, the cloned network can accurately integrate angular velocity. Note that, as in previous models [70], the gain of integration can deviate from 1 at different points along the ring due to the underlying geometry of the population dynamics (Fig. 4 S1).

**Figure 5:**
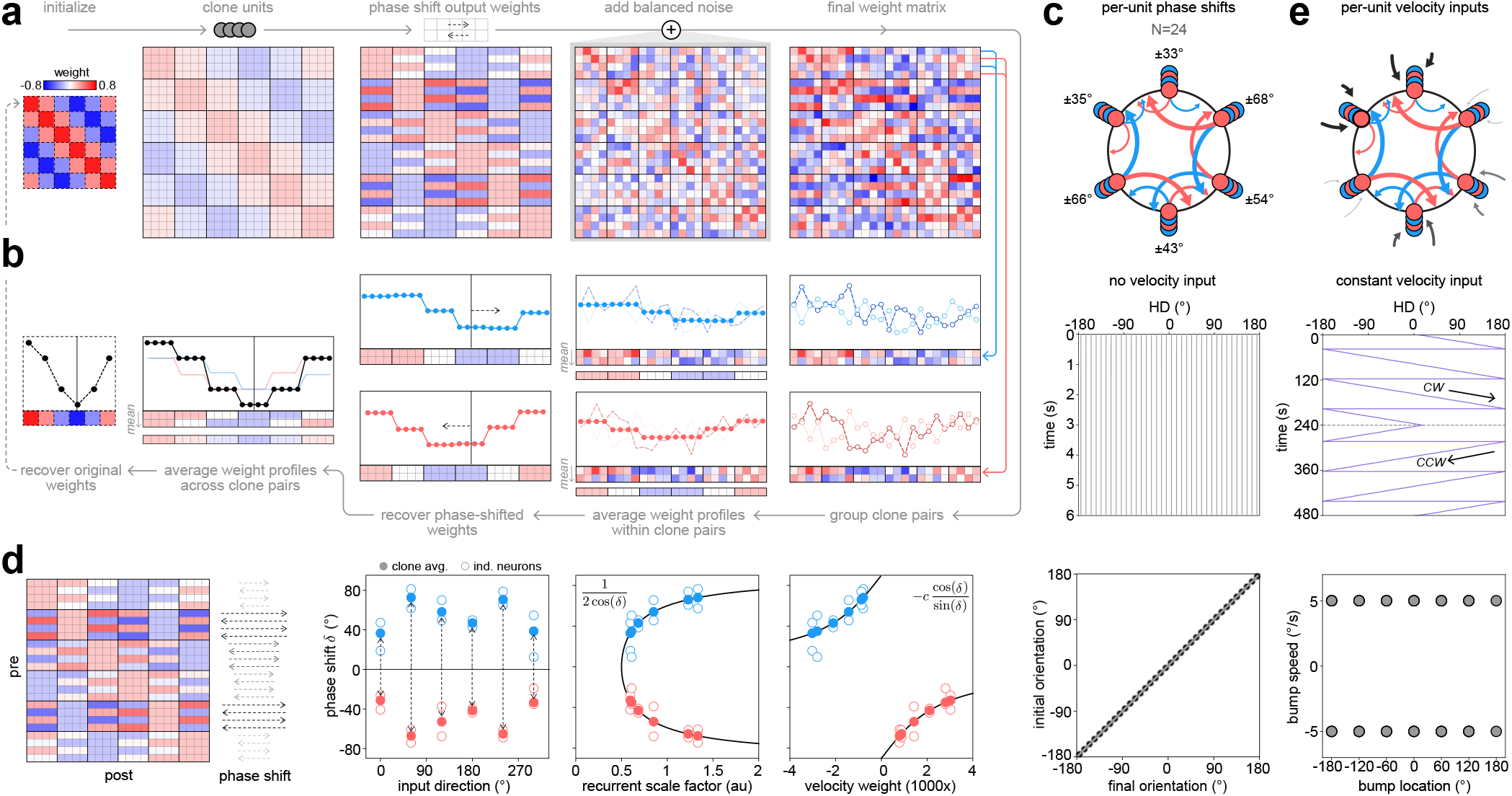
Constructing heterogeneous networks with balanced noise. **a)** As before, we begin with a small continuous attractor network (as in Fig. 4a; first column), and clone each unit four times (second column). Within each group of clones, we create two pairs of balanced phase shifts (third column); and add noise that is balanced across each pair (fourth column). The underlying hidden symmetries are completely masked in the final weight matrix (fifth column). **b)** Recovering the hidden symmetries in the network requires knowledge of the set of clones, and the pairs of neurons with balanced phase shifts (fifth column). Averaging the weights across each pair (fourth column) recovers the original phase shifts (third column). The average of these two phase-shifted weight profiles (second column) then reveals the original connectivity profile (first column). **c)** Even in the presence of additive noise in the recurrent weights, the cloned network maintains a bump of activity that can persist at any orientation in the absence of input. **d)** When phase shifts (first panel) are obscured by additive noise, velocity scaling rules must be applied based on the average phase shifts within each clone pair (filled markers, second through fourth panels). Additive noise alters the per-unit phase shifts, which in turn leads to inaccurate scaling of recurrent and velocity input weights (open markers, second through fourth panels). **e)** When recurrent and velocity input weights are appropriately scaled (based on the average phase shift within clone pairs, as highlighted in panel d), the cloned network can accurately integrate angular velocity.

**Figure 6:**
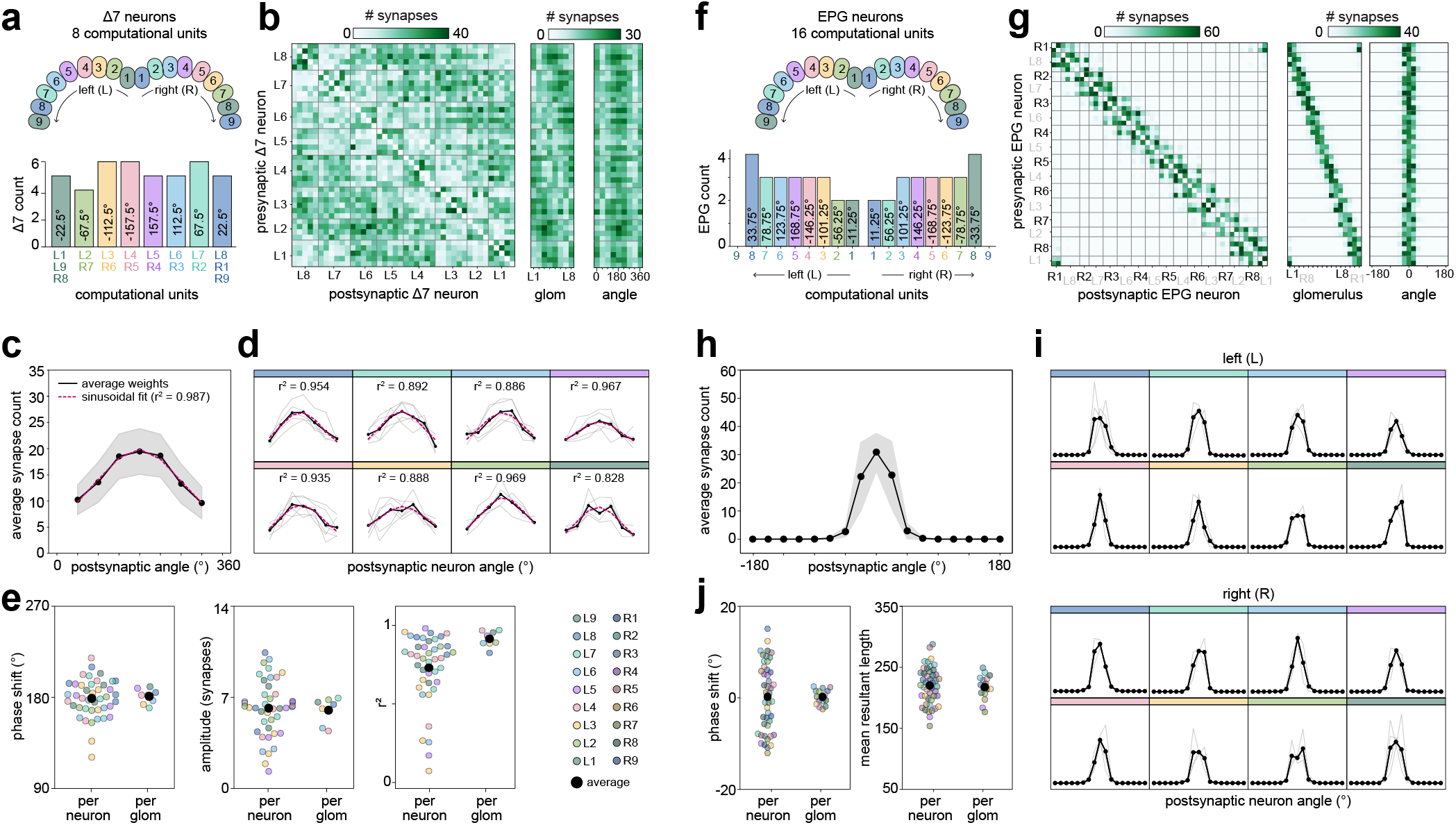
Recovering hidden symmetries in the fly connectome (CNS volume). **a)** Top: PB glomeruli colored by preferred tuning direction inferred from physiological measurements [68] and network modeling (Fig. 2h). Bottom: Δ7 neurons can be assigned to one of eight computational units that contain putative dynamical clones that share the same tuning direction (angles shown inside each bar). See Fig. 6 S2 for morphological renderings of all Δ7 neurons. **b)** Left: neuron-to-neuron connectivity matrix for all Δ7 neurons, grouped by computational unit (labeled L1 to L8). Notice that the connectivity is approximately symmetric but with considerable heterogeneity. Middle: connectivity from individual Δ7 neurons onto computational units, obtained by averaging across postsynaptic neurons (columns) that share a computational unit in the left panel. Right: phase-aligned connectivity, obtained by circularly shifting each row in the middle panel to align the connectivity profiles for direct comparison. **c)** Average axo-dendritic connectivity profile, obtained by averaging the phase-aligned profiles in the right panel of b (excluding the axo-axonic connections at the 0-degree and 360-degree columns). Black line marks the average weight profile (gray regions show mean ± standard deviation), with the sinusoidal fit shown in red (*r*^2^ = 0.99). **d)** Same as in (c) but broken out by groups of putative dynamical clones. Gray lines are individual Δ7 neuron connectivity profiles (average is shown in black), along with the sinusoidal fit (red line). **e)** Sinusoidal fits to individual Δ7 neuron weight profiles (gray lines in d) and clone-averaged profiles (black lines in d) were used to extract and compare the connectivity phase shifts (left panel), amplitudes (middle panel), and *r*^2^ values (right panel). Notice that the clone-averaged weight profile is more sinusoidal than the profiles of nearly all individual neurons, consistent with the presence of hidden symmetries. **f-j)** Same as in a-e but for EPG neurons. Note that the EPG connectivity profile is not sinusoidal, so phase shifts and connectivity amplitudes were estimated using a population vector average (see Methods).

### A machine-learning-based exploration of heterogeneity in ring attractors

Are there novel attractor architectures that could explain the heterogeneous wiring observed in the fly connectome? To explore the solution space of candidate attractor architectures, we trained RNNs to track head direction (HD) by integrating angular velocity (Fig. 2a; [56–61]). We considered three classes of RNNs whose network weights ranged from fully ring-constrained, with symmetric connectivity as in classical models (Fig. 1c), to fully unconstrained, with connectivity that lacks strict symmetry requirements and could thus permit architectures with heterogeneous connectivity. A third class of connectome-constrained networks [62, 63] allowed us to place the fly’s attractor architecture in this broader solution space.

All three network classes learn to track heading, but they do so using different internal mechanisms. Dissecting these mechanisms required us to generalize some of the analyses that are typically used to characterize units in these networks, which we describe here before detailing network-specific results. Classical models are hand-designed using rings of neurons that share the same structural and functional properties. As a result, neurons within each ring can naturally be grouped according to their connectivity patterns and response properties, and individual rings are easy to interpret, as in Fig. 1c. In contrast, our RNN formulation is more general and allows individual units to develop unique connectivity patterns and response profiles, especially in networks with unconstrained weights that allow for independent phase shifts across units (Fig. 2b). As a result, units cannot always be grouped into distinct structural or functional rings, and thus are more difficult to interpret. To overcome this challenge, we first computed single-unit tuning curves that characterize how the firing rate of each neuron depends on the network’s input (angular velocity) and output (HD; Fig. 2c). We then computed each unit’s preferred HD, which specifies a unit’s “input direction” and which can be used to order units by phase along a global topological ring (Fig. 2d). For each unit along this ring, we then computed the weighted average HD of the neurons to which it projects, which specifies the unit’s “output direction” (note that this relies on both structural and dynamical properties of individual units in the network). The difference between a unit’s input and output direction is then a measure of its phase shift (Fig. 2e, Methods). These quantities allow us to compare and contrast the solutions learned by each network class, revealing a large space of architectures with functional and structural features that depend on the imposed constraints.

Ring-constrained networks are built from three rings of neurons that each receive a shared velocity input (Fig. 2f, left). As in classical models (Fig. 1c), connections within and between rings are constrained to be circularly-symmetric versions of a shared connectivity profile; however, unlike classical models, each such connectivity profile can vary in its shape and phase-shift (Methods). As a result, these networks learn a variety of solutions that approximate both classical models and novel variants (shown in Fig. 2 S1-S4). For example, these networks learn approximations to canonical 3-ring models (Fig. 2f, left), with tuning curves that can be clustered into 3 rings (Fig. 2g, left). Consistent with classical models [8], units in each ring evenly tile the space of preferred head directions (Fig. 2h, left), and rings with phase-shifted connectivity receive stronger velocity input (Fig. 2i, left).

Connectome-constrained networks are composed of units whose connectivity is constrained to approximate that of individual neurons in the fly connectome (Fig. 2f, middle). In agreement with previous physiological recordings [24, 38, 43, 44], these networks learn tuning curves that fall into 5 functional clusters (Fig. 2g, middle): one cluster that encodes “pure” head direction (with units that correspond to EPG, PEG, and Δ7 cell types), and four clusters that conjunctively encode head direction and angular velocity (with units that correspond to the left and right PENa and PENb cell types). In agreement with previous light-level and connectome-based anatomical analyses [38, 39, 54], the conjunctively-tuned units have clockwise and counterclockwise phase shifts, but with variability across units (Fig. 2i, middle). Importantly, unlike the evenly-spaced HD tiling observed in classical models, the units in connectome-constrained networks cluster around distinct orientations (Fig. 2h, middle). In particular, units corresponding to Δ7, left EPG, and right EPG neurons each cluster around 8 distinct orientations, with the Δ7 population preferring orientations that are approximately halfway between the left and right EPG populations, consistent with previous functional measurements [44, 68].

Unconstrained networks are composed of units that can each learn their own phase-shifted connectivity profile and velocity input strength [58, 59]. As a result, these networks learn architectures with highly heterogeneous connectivity (Fig. 2f, right). The resulting tuning curves span a continuum and lack the clear functional clusters observed in ring and connectome-constrained networks. (Fig. 2g, right). Moreover, unconstrained networks have heterogeneous phase shifts and velocity input strengths, and units in these networks cannot be assigned to distinct rings based on these properties (Fig. 2i, right). These features deviate substantially from the strict symmetry and modularity that define classical models. In addition, individual units cluster around distinct orientations (Fig. 2h, right), a pattern that resembles the clustering observed in connectome-constrained networks.

These results reveal a large space of architectures that vary in their degree of modularity and heterogeneity (Fig. 3). At one extreme, ring-constrained networks learn crystalline architectures in which groups of units can be assigned to distinct rings that share the same velocity tuning and phase shifts; within each ring, individual units evenly tile head direction. At the other extreme, unconstrained networks learn heterogeneous architectures in which individual units vary substantially in their velocity tuning and phase shifts and thus cannot easily be grouped into distinct rings; however, units can be clustered based on their tuning to head direction. Connectome-constrained networks share properties of both extremes: units can be grouped into rings that share similar velocity tuning and phase shifts; within each ring, individual units vary in their velocity tuning and phase shifts but can be clustered based on their tuning to head direction.

When taken together, these results hint at coordinated changes in network parameters that leave performance roughly unchanged. Such coordinated changes would lie along “sloppy” dimensions in parameter space [69]; that is, directions along which connectivity can vary without degrading attractor dynamics or velocity integration. This is in contrast to “stiff” dimensions along which even small variations in connectivity substantially degrade performance (as in Figure 1g-h). If the heterogeneous connectivity observed in our trained RNNs reflects an underlying sloppy dimension in parameter space, it would preserve rather than destroy attractor dynamics. Moreover, because this heterogeneity always emerges alongside clustering in HD tuning, this redundancy in HD tuning may provide the mechanism for creating these sloppy dimensions, an idea we investigate next using theory.

### Dynamical clones enable heterogeneous connectivity in continuous attractors

Our RNNs discovered three key features that are present in connectome-constrained and unconstrained networks but are not found in canonical models: heterogeneous phase shifts, heterogeneous velocity tuning, and clustered head direction tuning. We sought to extend classical models to include these features without sacrificing exact continuous attractor dynamics. To this end, we developed an algorithm (Figures 4 and 5) that alters the structural connectivity of classical attractor networks [40] while preserving their functional performance, building in sloppy dimensions in the process. The algorithm identifies a set of “equivalence operations” that transforms a symmetric connectivity matrix into a heterogeneous one while maintaining exact continuous attractor dynamics. Crucially, by understanding how these operations generate hidden symmetries, we also specify how to reverse the algorithm to recover hidden symmetries from heterogeneous networks, an approach we subsequently apply to the fly connectome.

The algorithm’s core equivalence operations replace individual neurons with a redundant and dynamically equivalent set of units that share the same input direction but can differ in their output direction. We refer to these redundant units as “dynamical clones” because they are constructed to share the same firing rate in the absence of external velocity input. We require that each set of clones collectively preserves the summed output onto every postsynaptic neuron, which guarantees that in the absence of external input, each larger network is functionally equivalent to its starting network. The number of dynamic clones in a set can be arbitrarily large, and different sets can contain different numbers of dynamical clones; for simplicity, we illustrate the core algorithmic operations using small networks with a fixed number of dynamical clones per set. These algorithmic operations allow us to alter connectivity in three distinct ways, as illustrated in Fig. 4a (see Methods). First, the net self-coupling between dynamical clones must preserve the self-coupling of the original neuron; this weak constraint enables connectivity patterns that have no self-connections. Second, dynamical clones can support balanced additive noise: if one unit synapses onto a postsynaptic neuron too strongly, another clone in the set can weaken its connection onto the same postsynaptic neuron, such that their summed output is preserved. Third, each set of clones can support arbitrary phase shifts onto the rest of the network, provided that these phase shifts are counterbalanced and appropriately scaled across the set of clones. When combined with an appropriate velocity input weight, this enables accurate velocity integration. Together, these “equivalence operations” allow us to transform idealized networks with perfect symmetries [40] into functionally equivalent networks with hidden symmetries (see Methods and SI Materials Section 4 for a detailed derivation).

These hidden symmetries are not readily apparent in recurrent weights, but they nevertheless ensure that each network can support an accurate and persistent activity bump in the absence of velocity input (Fig. 4b). Accurate path integration further requires that the bump of population activity move at the same speed as the input velocity; in other words, the “gain” of integration must be one. In classical 3-ring models, this gain is determined by the magnitude of the phase-shift and the strength of the projections from the side rings back to the center ring; projections with larger phase shifts and stronger weights are more effective at moving the bump than those with smaller phase shifts and weaker weights ([9, 10]; SI Materials Sections 2.3 to 2.5). We used this to derive scaling operations for individual units in our cloned networks (Fig. 4c): if a unit gains a phase shift *δ*, it must strengthen its recurrent weights by a factor 1/(2 cos(*δ*)), and reduce its input velocity weight by a factor cos(*δ*)/ sin(*δ*) (see Methods and SI Materials Section 4.4.3).

The symmetries in these networks can be further masked by balanced noise (Fig. 5). To illustrate this, we constructed a network with four dynamical clones per input direction (see Methods). In the absence of any additional phase shifts, the additive noise need only balance across the set of four clones; however, when we introduce phase shifts into the network, the noise must balance in a way that preserves the net phase shift within each group of clones. For example, consider pairs of phase shifts in which two clones share a clockwise phase shift and two share a counterclockwise phase shift (Fig. 5a). When noise is appropriately balanced across each pair of clones, both the phase shifts and the additive noise cancel across the set of four clones (Fig. 5b). Empirically, we find that this noise can be of the same magnitude as the weights themselves while still supporting stable population dynamics, and can thus substantially mask the symmetric connectivity (far right panel of Fig. 5a). Despite this substantial heterogeneity, the resulting network nevertheless maintains a persistent activity bump at any orientation (Fig. 5c). To move this bump, the same velocity scaling rules still apply (Fig. 5d, filled markers), which enables accurate integration with uniform gain (Fig. 5e). This example highlights an important issue when attempting to recover hidden symmetries in heterogeneous networks like the connectome: if phase shifts are masked by noise, they are no longer apparent at the level of individual units. As a result, directly inferring the per-unit phase shift from the noisy weight profile leads to erroneous predictions of the recurrent and velocity input weights (Fig. 5d, open markers). Instead, recovering the true phase shift requires knowledge of the units that collectively operate as dynamical clones.

When taken together, these results show how a network gains extra degrees of freedom once multiple units share the same input direction. One clone can shift its output slightly clockwise and another slightly counterclockwise, so long as their summed output preserves the original connectivity profile. This constitutes a core sloppy dimension in our construction: larger phase shifts require larger recurrent weights to reconstruct the same total output. In the absence of any additional scaling, these shifted and scaled recurrent weights would move the bump too quickly; by reducing the corresponding velocity input weights, the network can recover the appropriate gain of integration. The result is a coordinated family of parameter changes — larger phase shifts, larger recurrent weights, and weaker velocity input — that leaves the overall computation unchanged. Unlike classical 3-ring models, these sloppy dimensions operate on pairs of dynamical clones instead of on distinct rings (as schematized in Fig. 3), allowing for a larger space of weight solutions. Balanced noise can further broaden these sloppy dimensions by allowing one clone with overly strong outputs to be balanced by a clone with weaker outputs.

Although we illustrated these sloppy dimensions for single network instantiations, this construction algorithm can generate an infinite space of heterogeneous ring attractor architectures that are dynamically equivalent to classical symmetric architectures. Individual units can be cloned an arbitrary number of times, with arbitrary phase shifts, and different groups can have a different number of clones. These clones allow the introduction of phase-shifted connectivity and balanced noise that create hidden symmetries that are not visually apparent in the recurrent weights. When combined with appropriately scaled velocity inputs, these networks smoothly integrate angular velocity, just as in classical models. Finally, with knowledge of the neurons that function as dynamical clones, it is possible to reverse this algorithm to reveal hidden symmetries, a finding we use to revisit the heterogeneous connectivity of the fly’s neural compass.

### Hidden symmetries in the fly’s compass network

The cloning algorithm described above shows how heterogeneous connectivity can hide symmetries that are indicative of a continuous attractor. It also suggests that reversing the algorithm can reveal such hidden symmetries, if they are present, and it makes two core predictions about the signatures of those symmetries. First, there should be multiple units that are tuned to the same preferred direction and that operate as dynamical clones. Second, noise should be balanced across clones, such that averaging their weights reveals the hidden symmetry required for continuous attractor dynamics. Importantly, weight averaging is only a method for revealing hidden symmetries if they exist; if they do exist, it suggests that the heterogeneous connectivity can — by itself, rather than as noise that must be averaged over — support continuous attractor dynamics (as shown in Fig. 4 and Fig. 5).

The fly compass network exhibits a key property that suggests its heterogeneous connectivity might reflect hidden symmetries: neurons within each of its five key cell types share tuning to particular head directions (Fig. 2h). These tunings are largely genetically predetermined and can be predicted based on innervation patterns within a region called the protocerebral bridge (PB; [39, 44, 54, 68]). The PB is composed of 18 glomeruli that each contain the dendrites or axons of neurons that share a similar directional tuning (Fig. 6a and 6f). Each neuron type is composed of many individual neurons – Δ7 (42), EPG (46), PEG (18), PENa (20), and PENb (22) – with anywhere from 1 to 6 neurons per glomerulus [39]. Physiological recordings [44, 68], connectivity analyses, and our RNN simulations (Fig. 2h) show that these 18 glomeruli can be reduced to 8 or 16 “computational units” by grouping neurons that share the same directional tuning. Together, the fact that tuning is shared by neurons within a computational unit suggests their potential to be dynamical clones; of the neuron types that display this shared tuning, the Δ7 and EPG neuron types are the most likely to contain dynamical clones since they have the largest number of neurons per computational unit (Fig. 2h). These two neuron types are also thought to ‘purely’ encode head direction via recurrent connectivity that is approximately circularly symmetric.

We began by studying the recurrent connectivity between Δ7 neurons (Fig. 6a-e) obtained from the recent Janelia CNS EM volume [35]. A previous connectome analysis suggested that Δ7 neurons have sinusoidal recurrent weights that could reformat EPG activity for downstream vector computations [39, 68]. However, because classical attractor models invoke sinusoidal connectivity, we reasoned that these recurrent Δ7 connections might support attractor dynamics as well. In addition, Δ7 neurons have nearly twice as many neurons per computational unit compared to the next most numerous type, the EPG neurons: Δ7 neurons have 42 neurons across 8 computational units, while EPG neurons have 46 neurons across 16 computational units. Fig. 6b shows the recurrent connectivity between Δ7 neurons, grouped into blocks of neurons that comprise the same computational unit and are likely to function as dynamical clones. Anatomically, each Δ7 neuron innervates all glomeruli in the PB, with axonal arbors in 2-3 glomeruli that share the same directional tuning and with dendritic arbors in the remaining glomeruli (Fig. 6a, inset; Fig. 6 S2). Interestingly, each Δ7 neuron receives the strongest dendritic input from Δ7 neurons with the opposite directional tuning. Since Δ7 neurons are thought to inhibit one another [38], this connectivity could implement the long-range inhibition required by classical models. In addition, Δ7 neurons that share a computational unit have axo-axonal synapses that could be excitatory or inhibitory, depending on the types of receptors expressed in the axonal compartment [38, 71]. Here, we focus on axo-dendritic synapses and ask whether the heterogeneous connectivity between Δ7 neurons could support their function as dynamical clones.

To this end, we reverse our cloning algorithm by first computing the average output from each Δ7 neuron onto all postsynaptic Δ7 neurons that share a computational unit (Fig. 6b, middle panel). Next, we shifted these weights to align their output profiles for direct comparison (Fig. 6b, right panel). In agreement with predictions from classical models, the average axo-dendritic output profile is well fit by a sinusoid, accounting for over 98% of the variance (Fig. 6c). Similarly, consistent with the form of heterogeneity introduced by our construction algorithm, the output profile of individual Δ7 neurons is less sinusoidal than the average output of the set of Δ7 neurons that function as putative dynamical clones (Fig. 6d-e). This suggests that a large fraction of the heterogeneity in per-neuron output weights comes in the form of balanced noise, where overly strong connections from one Δ7 neuron are balanced by weaker connections from another Δ7 neuron in the same computational unit. A similar analysis applied to EPG neurons and their 16 computational units gives similar results (Fig. 6f-i). Applying the same analysis to the hemibrain EM volume [33] also yielded similar results (Fig. 6 S1), indicating consistency across fly connectomes. Taken together, the finding that the aggregated weights of putative dynamical clones are more symmetric than their single-neuron counterparts is consistent with the presence of hidden symmetries in the fly compass network.

## DISCUSSION

Foundational theoretical models can explain how recurrent neural networks stably encode and update internal representations of continuous angular variables such as head direction (Fig. 1). Recent dense connectomes have provided strong support for the core architectures proposed by these models, while also revealing substantially more wiring heterogeneity than classical ring attractor models can tolerate. To understand this discrepancy, we studied relationships between structural connectivity and neural dynamics in recurrent networks trained to accurately path integrate (Fig. 2). By varying the structural constraints imposed on these networks, we identified several sources of heterogeneity that arise in both connectome-constrained and unconstrained networks, but are absent from existing theoretical models (Fig. 3). This heterogeneity hinted at the existence of sloppy dimensions that permit coordinated changes in network parameters without altering network performance. We formalized this idea by construction: by cloning the dynamics of individual neurons in small attractor networks, we introduced heterogeneous connectivity while preserving exact continuous attractor dynamics (Fig. 4 and Fig. 5). When combined with existing theory, this allowed us to precisely derive a set of sloppy dimensions and show that the space of continuous attractor architectures is larger than previously appreciated. Finally, by identifying putative dynamical clones in the fly’s compass network, we applied this construction principle in reverse to reveal hidden symmetries in connectivity that could support continuous attractor dynamics (Fig. 6).

### Combining machine-learning-based discovery with theory to understand biological circuits

Task-trained recurrent neural networks are widely used to generate hypotheses about how neural circuits implement specific computations ([56–61, 72–75]; reviewed in [76–79]). To maximize flexibility in the representations learned by these networks, many studies avoid imposing structural constraints, and focus their analysis on emergent population dynamics. As a result, the structural connectivity of trained networks is often disregarded. This tendency is reinforced by the fact that, in most biological systems, ground-truth connectivity is unavailable and therefore cannot be used as a constraint or as a point of comparison.

The fly compass system provides a unique opportunity to evaluate the utility of structural constraints in interpreting the dynamics of trained networks. Using an ensemble modeling approach [80, 81], we found that knowledge of structural connectivity—and specifically, the phase-shifted relationship between a unit’s preferred input and output direction—was critical for understanding how velocity is integrated within the network. This relationship only emerged when analyzing the precise connectivity of individual units, and would have been obscured by the weight averaging procedures that are sometimes used to analyze trained networks [58, 59]. These findings align with a growing body of work linking the structure of recurrent weights to the dynamics they produce, enabling an implementation-level understanding of network function and the extraction of computational motifs [23, 58, 59, 70, 82–87].

The precise structural constraints imposed on each network also impacted the mechanistic nature of the solution that was learned. Thus, accurate task performance alone is not sufficient to identify biological circuitry [60, 62, 63, 88, 89]. Instead, achieving implementation-level understanding of biological circuit function will likely require incorporating not only connectomic constraints, but also information about biophysics and neuromodulation [90–92]. Nevertheless, by interpreting trained networks through the lens of both theoretical models and the known organization of the fly compass, we found that all network solutions relied on balanced, phase-shifted connectivity to implement angular velocity integration. Thus, unconstrained RNNs can recover core aspects of biological implementation, and, when analyzed within appropriate theoretical frameworks, can help inspire new theory.

### Dynamical clones expand the space of continuous attractor architectures

Theoretical models necessarily abstract biological detail to gain analytical tractability. This abstraction has enabled deep functional insights, but it also creates a gap between idealized models and real biological circuits. In the fly compass system, this gap is especially apparent in the form of heterogeneous synaptic connectivity that, according to classical continuous attractor theory, should disrupt stable bump dynamics [40, 42, 50, 51, 55]. Rather than directly fitting this biological complexity into existing models, our dynamical clone algorithm starts from idealized ring attractor networks and builds in heterogeneous connectivity while preserving continuous dynamics. In the process, the construction algorithm brings theoretical models closer to biology without sacrificing analytical tractability. Importantly, this algorithm can be applied to any recurrent weight matrix in order to generate ensembles of network structures that preserve a given set of specified dynamics, regardless of which computations those dynamics support.

Dynamical clones enable architectural flexibility via a form of overparameterization. Additional units create sloppy dimensions: coordinated directions in parameter space along which connectivity can vary without changing the network’s performance [69]. The notion of sloppy dimensions is related to similar ideas across fields, including “flat minima” in the loss landscapes of machine learning models [93] and “degenerate” or “multiple” solutions in systems biology and neuroscience [90, 94]. In our case, degenerate solutions form a continuum in parameter space rather than existing as isolated points. For example, pairs of dynamical clones can arbitrarily shift their weight profiles, provided that their amplitudes and velocity input weights are rescaled appropriately to preserve continuous dynamics and accurate integration. Balanced weight noise enables further flexibility without degrading function. As a result, a single idealized network can generate entire families of functionally equivalent but structurally heterogeneous networks. Notably, while dynamical clones are constructed to preserve steady-state dynamics in the absence of input, the phase-shifted connectivity they enable allows the network to integrate angular velocity, demonstrating how additional sloppy dimensions can support new computational abilities.

These constructions also link naturally to existing theory. By building on recent theoretical work in small networks [40], we ensured that the resulting networks can generate continuous attractors that resemble classical 2- and 3-ring models [8, 10], as well as more unconstrained architectures (SI Materials Section 5), depending on whether the added clones share the same phase-shift (as in classical models) or not (as in unconstrained networks). More broadly, our results connect continuous attractor theory to the idea that overparameterization can support robustness. In machine learning, overparameterization is known to improve performance and can enable multiple solutions that span flat regions of the loss landscape, consistent with the existence of sloppy dimensions and degenerate solutions [93, 95–97]; in systems biology and neuroscience, overparameterization is thought to support degenerate solutions that enable robust dynamics despite external perturbations and internal variability in structural and dynamical properties [90, 94, 98, 99]. Our findings suggest that analogous principles can help mitigate the classical fine-tuning problem in continuous attractor networks by expanding the set of network architectures that preserve desired dynamics.

### Dynamical clones provide a path to structured yet robust biological wiring

Dynamical clones also suggest a concrete mechanism by which biological attractor circuits might tolerate heterogeneous connectivity. In architectures where neurons are uniformly distributed around a ring (Fig. 1a and g), weight noise at one synapse can only be compensated by changes at that same synapse, or by adding additional rings [42]. In contrast, when multiple neurons share the same heading tuning, their weights can collectively compensate for one another. For example, a network that is constrained to use 16 units could distribute those units evenly around a ring, as in classical models, or organize them as 4 groups of 4 clones, 8 groups of 2 clones, or any other partition. All of these configurations can produce exact continuous attractor dynamics within the same 16 × 16 weight space, but clone-based architectures permit more sloppiness in the specific instantiation of those weights.

This type of compensation could arise naturally from physical and developmental constraints. For example, if one neuron forms more synapses onto a target, its processes may occupy space that reduces the number of synapses formed by a clone with similar tuning. Such local competition would generate balanced weight noise and reduce variability relative to independently distributed synaptic weight noise. Our connectome analyses of the fly compass provide support for this type of balanced noise: the aggregated weights of putative dynamical clones are more symmetric than their single-neuron counterparts.

At the same time, the fly compass does not appear to exploit the full unconstrained space of possible clone-based architectures. Instead, our results suggest that it combines dynamical clones and heterogeneous weights within a modular organization. This modularity may also reflect developmental constraints. Because the fly compass is largely genetically hardwired, architectures that can be specified by fewer wiring rules may be easier to encode and implement through development [100, 101]. For example, a circularly-symmetric connectivity matrix requires at most *n* underlying weight parameters for an *n* × *n* weight matrix, whereas a fully unconstrained matrix could require up to *n*^2^ parameters. Similarly, because every PEN neuron shares the same approximate phase-shift magnitude, their velocity inputs can also be approximately uniform [39, 53] . In non-modular architectures with variable phase shifts, by contrast, velocity inputs would need to be individually calibrated, complicating the wiring problem and potentially requiring online learning. Such learning may occur during development or into adulthood [102], but the fly’s modular structure simplifies the problem considerably.

Several caveats remain. Synapse counts are only a proxy for true synaptic strengths, and our weight-averaging procedure could underestimate the symmetry of the underlying functional connectivity. In addition, variance in synapse detection, proofreading completeness, and other aspects of connectome reconstruction could obscure perfect symmetries even if they were present. Further, the biophysical properties of individual neurons could contribute to functional symmetry in the network in ways that we do not fully explore in our circuit-level analyses. Nevertheless, the fact that aggregated weights of putative dynamical clones are more symmetric than single-neuron weights supports the idea that apparent heterogeneity contains hidden symmetries that can support continuous attractor dynamics.

### Outlook

Our results provide a framework for leveraging machine-learning-based discovery in closed-loop with theoretical models and experimental measurements of connectivity and physiology to iteratively refine our understanding of circuit-level computations. Applied to the fly compass network, this framework shows how networks with heterogeneous connectivity can nevertheless produce continuous attractor dynamics, and suggests that such networks may require less fine-tuning than previously believed. More broadly, our approach offers a way to generalize idealized models of neural computation to the richer and messier architectures found in biological circuits, helping bridge the gap between theory and biology.

## ACKNOWLEDGEMENTS

We thank Kristin Branson for feedback on early versions of the connectome-constrained RNNs and helpful discussions; Marcella Noorman and Srini Turaga for helpful discussions; members of the Hermundstad and Jayaraman labs for feedback throughout the project; Marisa Dreher for help with neuron renderings; and Dan Turner-Evans, Cedric Allier, Chris Kymn, Chad Sauvola, Tzuhsuan Ma, and Josh Dudman for feedback on the manuscript. This work was funded by the Howard Hughes Medical Institute.

## METHODS

### RNNs trained to integrate angular velocity over time (Figure 2)

Head direction systems integrate an angular velocity signal over time to update an internal estimate of the animal’s allocentric heading. Similar to previous work [58, 59], we trained RNNs to generate dynamics that support this computation, but with different types of constraints placed on the network’s recurrent weights and velocity input weights, as described further below. The dynamics of these networks are all governed by the same nonlinear dynamical system equation that describes a firing rate model [103], which we simulated in discrete time using Euler integration:

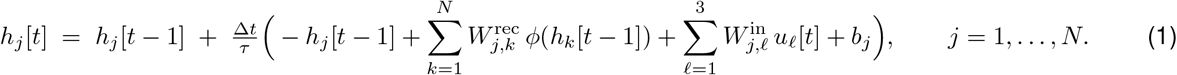

Where 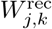 is the recurrent weight from presynaptic unit *k* to postsynaptic unit *j* (*N* = 156 units for all networks), *h*_*j*_ [*t*] is the subthreshold activity (voltage) of unit *j* at time step *t, ϕ*(*·*) is a sigmoid nonlinearity (*ϕ*(*h*) = 1/(1 + *e*^−*h*^)) that converts subthreshold activity into a firing rate, 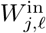 is the input weight conveying input channel ℓ to unit *j* (forming an *N ×* 3 input weight matrix), *u*[*t*] is the 3-dimensional input vector that carries a rotational velocity signal and the initial heading (described further below), *b*_*j*_ is the bias to unit *j, t* ∈ {1, 2, 3, … } indexes discrete steps, Δ*t* = 0.01 s (so physical time is *t* Δ*t*), and *τ* = 0.1 s.

Head direction is decoded from *ϕ*(*h*) using a linear readout layer:

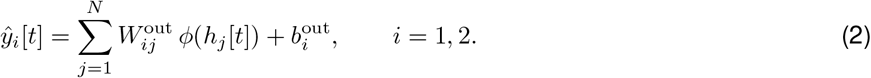

Where 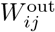 is the readout weight from unit *j* to output channel *i* (forming a 2 *× N* readout weight matrix), *b*^out^ is a 2 × 1 bias vector, and *ŷ*[*t*] is the model’s 2 × 1 output vector that should lie on the unit circle with a phase that matches the true head direction at time *t*. The trainable parameters for all networks are: *W*^in^, *W*^rec^, *b, W*^out^, and *b*^out^.

We use a mean-squared error loss function:

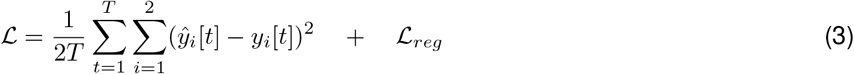

Where *y*[*t*] is the ground truth head direction, *θ*^hd^, encoded as *y*_1_[*t*] = cos(*θ*^hd^[*t*]) and *y*_2_[*t*] = sin(*θ*^hd^[*t*]). As described below, additional regularization terms (ℒ_*reg*_) were added to the MSE loss function to constrain the network’s recurrent weights depending on the type of network being trained. All networks were trained and simulated using PyTorch [104]. Code will be made publicly available upon publication.

#### Synthetic angular velocity generation

Similar to previous work [43, 58], training and test data were generated using an Ornstein-Uhlenbeck (OU) process to simulate a walking fly’s rotational velocity statistics. The discrete-time OU process is given by:

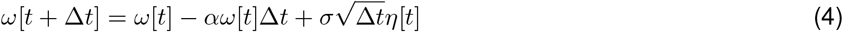

where *ω*[*t*] is the angular velocity at time *t, α* = 1*/τ*_*corr*_ is the inverse of the autocorrelation time constant, 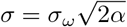 where *σ*_*ω*_ is the stationary standard deviation of the velocity, and *η*[*t*] ∼ 𝒩 (0, 1) is Gaussian noise. We used an autocorrelation time constant of *τ*_*corr*_ = 0.12 s and a velocity standard deviation of *σ*_*ω*_ = 40 deg/s.

To simulate periods where the fly stands still, we inserted pauses into the velocity traces. Pause durations were drawn from an exponential distribution with a mean of 2 s, capped at a maximum of 8 s. Pauses were inserted sequentially at random positions in the sequence until approximately 20% of the trial consisted of standing periods. During stops, head direction was held constant and the velocity input was 0. The ground truth head direction at each time step was computed by numerically integrating the velocity signal:

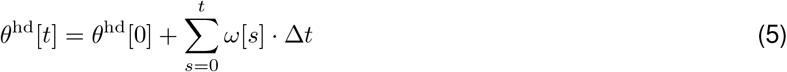

where *θ*^hd^[0] is a random initial heading drawn uniformly from [0, 360) degrees and *ω*[0] = 0. The input to the network, *u*[*t*], was a 3-dimensional vector:

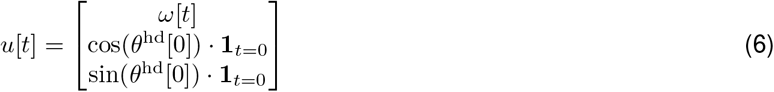

where the first component contains the angular velocity in degrees per second, and the second and third components encode the initial head direction as cos(*θ*^hd^[0]) and sin(*θ*^hd^[0]), respectively, at the first time step only (set to zero for subsequent time steps). This initial heading cue allows the network to learn to initialize the bump at different positions along the ring and integrate angular velocity to update the bump’s position thereafter.

#### Network training

The training data contained 200,000 simulate velocity trials that were divided into 2,000 batches of 100 trials each. Each trial consisted of 100 time steps with Δ*t* = 0.01 s, corresponding to 1 s of simulated time per trial. The networks were trained for 10 epochs, with a learning rate of 0.005 that decreased by a factor of 10 at epoch 5. The gradient was computed using the backpropagation through time (BPTT) algorithm [105], and the parameters were optimized using Adam [106]. Network-specific hyperparameters are listed in the sections below.

#### Network architectures

We trained three classes of networks with different parameter constraints, as described below. For each class, we trained 10 RNNs to ensure consistent results and to sample the types of solutions learned across networks (Figure 2 S1 to S4).

<u>Unconstrained RNNs</u> had no constraints placed on any of their trainable parameters, as is typical in field of task-trained RNNs. Each element of *W*^rec^ and *W*^in^ was independently initialized from 𝒩 (0, 1*/d*^2^), with *d* = 100. The recurrent units were all initialized with a bias of 1. *W*^out^ and *b*^out^ were initialized using PyTorch’s default Kaiming initialization. After initialization, all parameters were trained without constraints, allowing individual units to learn their own velocity input strengths (elements of 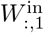) and phase-shifted connectivity patterns. These networks consisted of *N* = 156 recurrent units to match the total unit count in connectome-constrained networks.

<u>Ring-constrained RNNs</u> were inspired by classical three-ring attractor models and were constructed from three interacting rings of units. Each ring contained 52 units, for a total of 156 units. The 156 × 156 recurrent weight matrix was composed of 9 ring-to-ring submatrices indexed by *o* ∈ {1, 2, …, 9}. Each 52×52 submatrix *W*^(*o*)^ was a circulant matrix, where each row is a shifted version of its neighboring row (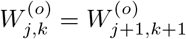 for all *j, k*, with indices modulo 52). This structure means that each submatrix is fully determined by a single learnable base vector *w*^(*o*)^ ∈ ℝ^52^, reducing the number of trainable recurrent weight parameters from 156^2^ = 24,336 to 52 × 9 = 468. Each base vector was initialized to produce a diagonal submatrix with diagonal elements set to 0.01. To favor smooth (approximately cosine-like) base vectors, we applied a smoothness regularization using a circular second-difference penalty:

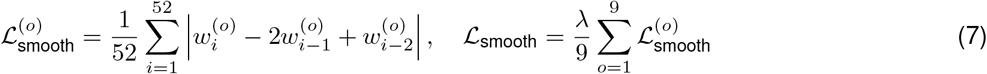

where 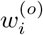 are the elements of the base vectors, indices are taken modulo 52 to enforce circular boundary conditions, and *λ* = 0.4 is the regularization strength. Importantly, even though base vectors were encouraged to be smooth, the angular phase shift was not constrained, allowing the network to learn novel phase-shifted connectivity patterns between rings. As in classical 3-ring models, units belonging to the same ring received the same angular velocity input strength, so the 156 velocity input weights in 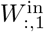 were populated using three learnable scalars (one for each ring) that were each initialized randomly from 𝒩 (0, 1*/d*^2^), with *d* = 100. Each recurrent unit’s bias was initialized to be 1. *W*^out^ and *b*^out^ were initialized as in unconstrained networks.

<u>Connectome-constrained RNNs</u> have recurrent weights that are constrained by the connectivity of the fly compass network [62, 63, 88], as described below. We include all HD-tuned neuron types that physiological recordings and perturbation experiments suggest are required for stable activity bumps and/or angular velocity integration: EPG (including EPGt), PEG, PENa, PENb, and Delta7. Each of these populations supports activity bumps that track the fly’s HD. Physiological recordings indicate that EPG, PEG, and Delta7 are primarily HD-tuned [24, 38], while PENa and PENb are conjunctively tuned to HD and angular velocity [43, 44]. Perturbation experiments have shown that silencing EPG, Delta7, PENa, or PENb neurons impairs the network’s ability to maintain stable activity bumps and integrate angular velocity [38, 44], suggesting that all 5 types are required for accurate compass dynamics. PEG silencing experiments have not been reported, but their main outputs are to PENb neurons whose silencing does impair bump dynamics [38, 44]. We also include an inhibitory neuron type known as ER6 that is thought to provide broad feedback inhibition to HD-tuned neurons [107], though we note that the compass network contains other potential sources of feedback inhibition that we do not attempt to capture. Similarly, we exclude neurons types whose silencing does not impact bump stability or angular velocity integration, including HD-tuned EL neurons [108], dopaminergic ExR2 neurons [109], and mechanosensory [110] or visually-responsive ring neurons [111–117], which are all involved in anchoring the compass to visual cues, a computation we do not study here. In performing connectome-constrained simulations, we do not aim to capture the full biological complexity of the fly circuit or make predictions for every function or computation that each neuron type might support. There are far too many unknowns to do so accurately – the actual network is composed of spiking neurons with multiple axonal and dendritic arbors that express a large number of neurotransmitter receptor types, and biological complexities like these could contribute to attractor dynamics in important but poorly understood ways. Instead, our connectome-constrained RNNs are a useful abstraction for studying how the fly compass network might support angular velocity integration through phase-shifted connectivity between populations of neurons with a mixture of HD and velocity tuning.

For connectome-constrained RNNs, the recurrent weight matrix contains 156 units and was initialized and regularized using a processed version of the raw synaptic connectivity matrix obtained from the Janelia CNS volume [35] using the neuPrint API [118]. We obtained the raw synaptic connectivity matrix by: (i) retrieving all neuron-to-neuron connections between the neuron types described above, (ii) excluding EPG synapses outside the PB and EB (iii) removing synaptic connections with fewer than 5 synapses, and (iv) assigning Delta7 and ER6 to be inhibitory and all other types to be excitatory. Next, we grouped type-to-type connections into three classes. For each type-to-type submatrix, we first computed the connection density as the proportion of non-zero connections, where a value of 1 reflects all-to-all connectivity and a value of 0 reflects no connectivity. We then used simple heuristics to classify submatrices into three classes: all-to-all, circulant, or sparse. Submatrices with connection density above 0.5 were treated as all-to-all (excluding submatrices involving Delta7) and were replaced by a constant matrix whose entries equal the average connection strength. Type-to-type submatrices between columnar types with connection density above 0.07 were treated as circulant. Submatrices that were neither all-to-all nor columnar were treated as sparse and all weights were set to zero. Lastly, we applied a “symmetrization” procedure to the type-to-type submatrices classified as circulant to generate symmetric connectivity as follows. First, we reduced each neuron-to-neuron connectivity submatrix to a glomerulus-to-glomerulus matrix by grouping neurons that innervate the same glomerulus and computed the average weight across all pairs of glomeruli. Next, we shifted each pre-synaptic row in the glomerulus-to-glomerulus matrix to align the weight profiles according to the relative phase difference between glomeruli and calculated an average weight profile. Lastly, this average weight profile was used to construct a circulant version of the glomerulus-to-glomerulus connectivity matrix (as in ring-constrained networks) which was then used to reconstruct a symmetrized version of the neuron-to-neuron connectivity matrix.

Let *W*^con^ denote the processed weight matrix of synaptic counts between pre- and postsynaptic units. The recurrent weight matrix was parameterized as:

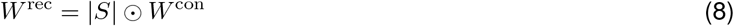

where *S* is a learnable per-connection scaling matrix, ⊙ denotes elementwise multiplication, and |*S*| ensures nonnegativity. This parameterization fixes the sign and sparsity of the connectome while allowing per-connection rescaling. Elements of *S* were initialized to be 0.01 for non-zero connections; elements where connections were absent were initialized to be 0 and kept their by masking their gradients during training. As in the fly circuit [53], velocity input was only provided to the PENa and PENb populations, with equal magnitude but opposite signs for the left and right populations. That is, similar to ring-constrained networks, the velocity input weights in 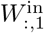 were populated from independently learnable scalars for the left and right PENa and PENb populations, with the rest of the elements set to 0 so that the other neuron types did not receive direct velocity input. The elements of 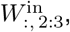, which set the initial bump position, were initialized as in ring-constrained and unconstrained networks. The recurrent bias terms were all initialized to 1. *W*^out^ and *b*^out^ were initialized as in unconstrained and ring-constrained networks.

A cosine distance regularization term was added to the loss function of connectome-constrained networks to keep the learned weights close to the connectome pattern, computed separately for each type-to-type connectivity submatrix:

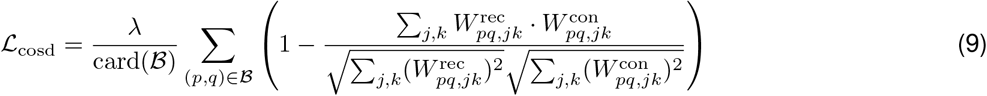

where 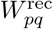 is the learned weight submatrix from presynaptic type *p* to postsynaptic type *q*, 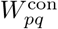 is the corresponding connectome-derived submatrix, indices *j* and *k* run over all unit pairs within each submatrix, ℬ is the set of all presynaptic-postsynaptic type pairs (*p, q*) for which the connectome-derived submatrix is nonzero, card(ℬ) is the number of elements in the set (cardinality), and *λ* = 1.0. This regularization allowed each type-to-type connectivity submatrix to scale the overall magnitude of its weights relative to other submatrices while restricting each submatrix’s connectivity structure to remain similar to the connectome. This was motivated by the observation that synapse counts, as obtained from the connectome, are likely to translate to different synaptic strengths depending on the presynaptic and postsynaptic unit types involved. Because cosine distance is scale-invariant, we additionally imposed a soft lower bound on the mean absolute weight magnitude for each type-to-type submatrix to prevent their connection strength from being scaled down below functional relevance (using a mean absolute weight threshold of *κ* = 0.05):

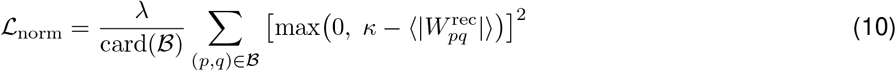

### Reverse engineering trained RNNs (Figure 2)

To characterize the computational mechanisms learned by each network, we computed tuning curves and connectivity-derived phase shifts for individual units.

#### Computing tuning curves

We started by computing a two-dimensional tuning curve for each unit by calculating its average firing rate as a function of the decoded head direction output and angular velocity input. To have enough data for averaging, we generated long test trials (12,000,000 time steps, corresponding to 120,000 s) using the same OU process described above and passed them through the trained networks. The network’s decoded head direction was computed from its output using the four-quadrant inverse tangent:

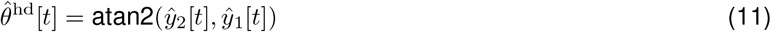

For each unit *k*, a 2D tuning curve was computed by averaging the unit’s firing rate in discretized bins of decoded head direction and angular velocity. We denote the binned head direction values as hd_*l*_ (*l* = 1, …, 128; uniformly spaced between ±180 deg) and the binned angular velocity values as av_*m*_ (*m* = 1, …, 128; uniformly spaced between ±80 deg/s), where the integer indices *l* and *m* index bins of the continuous variables *θ*^hd^ and *ω*, respectively. The 2D tuning curve is then:

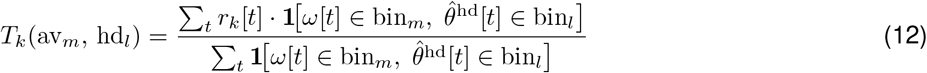

where *r*_*k*_[*t*] = *ϕ*(*h*_*k*_[*t*]) is the firing rate of unit *k* at time *t* and 1[*·*] is an indicator function that equals 1 when the velocity and decoded head direction at time *t* fall within the respective bins. From these 2D tuning curves, we computed marginal tuning curves for HD 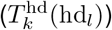 and AV 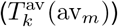 by averaging across the angular velocity or HD dimensions, respectively. The HD tuning strength of each unit was quantified as the max-min difference of its HD tuning curve. AV tuning strength was quantified as the median slope of the AV tuning curve.

#### Computing phase shifts

The preferred firing direction of each unit, also referred to as its input direction, was computed using a population vector average (PVA) by taking the circular mean of HD bins weighted by the unit’s firing rate in each bin:

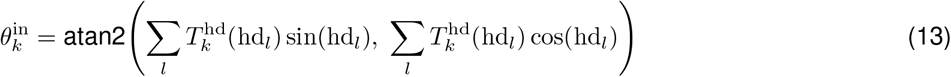

A unit’s input direction can be interpreted as its position on the global topological ring of units and reflects the angular position of the bump that causes the unit to fire at its highest rate. A unit’s “output direction” can be defined as the PVA of the preferred firing directions of all postsynaptic units it projects to, where the weights are the recurrent connection strengths onto postsynaptic unit *j* from presynaptic unit *k*:

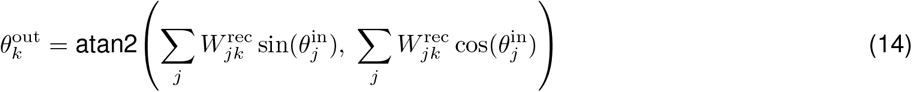

A unit’s output direction captures the location on the global ring to which it projects the strongest. The phase shift of each unit was then defined as the difference between its output and input directions:

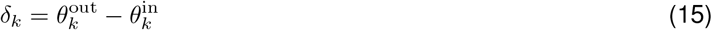

wrapped to the range [−180^°^, 180^°^). For example, a unit may have an input direction of 0-degrees (e.g. north) and project most strongly to units with preferred firing directions at 90-degrees (e.g. east), so its phase shift would be 90-degrees.

### Dynamical clone algorithm (Figures 4 and 5)

We developed an algorithm that transforms simple ring attractor networks with idealized connectivity into larger networks with heterogeneous connectivity while preserving continuous attractor dynamics. The algorithm operates through “equivalence operations” that clone the dynamics of individual units while altering their connectivity. In the sections below we provide a high-level overview of the algorithm as it relates to the results in Figures 4 and 5. A full derivation of the algorithm and its extensions can be found in SI Materials (Section 4).

Throughout this section we follow the clone indexing convention of the SI Materials: lowercase Latin subscripts (*i, j, k*) index units in the original (uncloned) network, and Greek subscripts (*µ, ν*) are suffixed to Latin indices to identify individual clones within a clone group. Thus *pµ* denotes clone *µ* of parent neuron *p*, and the set of all clones of neuron *p* is written 𝒞(*p*) = {*p*1, *p*2, …, *pm*}. Weights in the cloned network are written 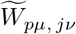 (from clone *jν* to clone *pµ*).

#### Starting ring attractor model

The algorithm can operate on any recurrent weight matrix, but for our application, we begin with a small threshold-linear ring attractor network of *N* units, as previously described [40]. Briefly, each unit’s preferred firing direction is:

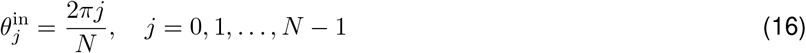

The connection strength between presynaptic unit *k* and postsynaptic unit *j* depends on the difference in their preferred firing directions according to:

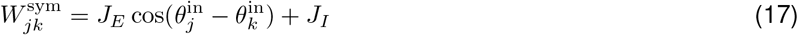

where *J*_*E*_ controls the strength of the cosine connectivity, *J*_*I*_ controls the strength of global inhibition, and 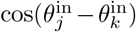 ensures circulant connectivity that is shift-invariant around the ring. For the starting networks used in Figures 4 and 5, we used *N* = 6, *J*_*E*_ = 4, and *J*_*I*_ = −0.5. This choice of *J*_*E*_ guarantees the network generates continuous ring attractor dynamics [40]. For visualization purposes, the DC component was removed from the matrices displayed in Figures 4 and 5.

For velocity integration, the model uses an asymmetric connectivity matrix defined as:

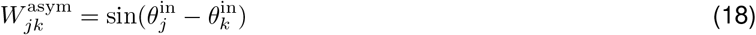

Similar to the RNN formulation, the dynamics of each postsynaptic unit *h*_*j*_ was governed by:

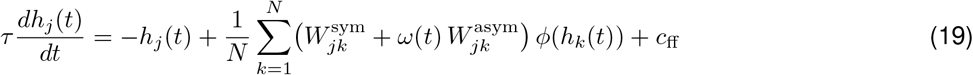

where *c*_ff_ = 1 is a constant feedforward input to all neurons in the network, *ϕ*(*·*) is a threshold-linear (ReLU) nonlinearity that transforms activity into a firing rate, *ω*(*t*) is the angular velocity signal that scales the asymmetric connectivity matrix, and *τ* = 0.1 s is the neuronal time constant. We simulated these continuous time dynamics in discrete time using Euler integration, as described above. The orientation of the bump at each moment in time was computed as the phase of firing rate vector’s first Fourier mode.

#### Duplication with phase shifts (Figure 4)

The algorithm used in Figure 4 introduces dynamical clones by duplicating each unit in the network and adding phase-shifted connectivity. This transforms the 6 × 6 symmetric starting matrix into a 12 × 12 cloned matrix with heterogeneous phase shifts and hidden symmetries. As shown in Figure 4a, for a parent unit *p*, the duplication into clones *p*1 and *p*2 proceeds as follows:

##### Step 1

Row and column duplication: Each duplication event starts by inserting a duplicate row and column into the weight matrix immediately after position *p*, initially copying all weights to and from the original unit. Copying the input weights to each clone ensures that both *p*1 and *p*2 receive the same total input from other units in the network, so long as the self-coupling input is also preserved.

##### Step 2

Self coupling between clones: The dynamical clone algorithm allows for flexible coupling between clones. In the case of Figure 4, where we duplicate each unit, the only requirement is that each clone in a duplicate pair (e.g. clones *p*1 and *p*2) receives the same total input weight as the original unit’s (*p*) self-coupling weight. That is:

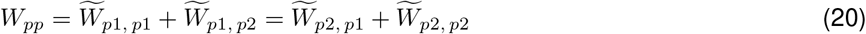

where *W*_*pp*_ is the self-coupling in the original network and 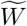 denotes weights in the cloned network. This flexible coupling between clones enables an infinite number of coupling patterns, including ones where units no longer have true self-coupling (i.e. 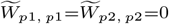), as in biological networks.

##### Step 3

Halve output weights: Next, to preserve the total output from the duplicate pair of clones onto all other units in the network, each clone’s output weights must be halved:

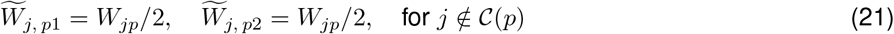

This ensures that all units that are postsynaptic to the dynamical clones receive the same total input that they received from the original unit so that their firing rate is preserved.

##### Step 4

Apply phase shifts with amplitude scaling: The key operation that allows us to introduce phase-shifted connectivity into a symmetric starting matrix is that a cosine connectivity profile can be split into two phase-shifted profiles whose sum equals the original profile using the trigonometric identity:

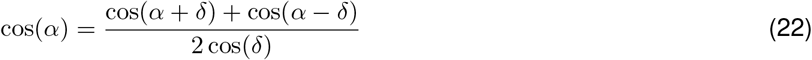

Using this identity, we shift the output weights from duplicate pairs in opposite directions (*δ*_*p*1_ = −*δ*_*p*2_), and scale the weights by 1/(2 cos(*δ*_*p*_)) to preserve the summed output onto each postsynaptic target:

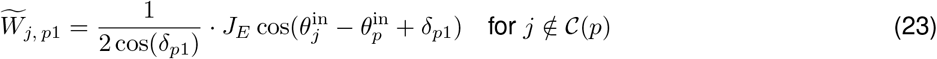

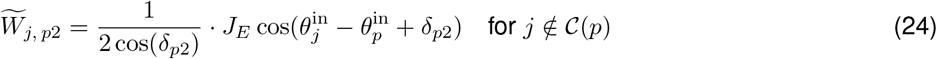

As shown in Figure 4, the weight scaling in steps 3 and 4 above can also be implemented by changing the slope of the unit’s non-linearity. These steps can be repeated to generate many clones per original unit but this requires keeping track of cumulative phase shifts and scaling factors. A simpler approach for generating multiple clones per original unit is described next.

#### Multi-clone model with balanced noise (Figure 5)

For generating networks with more than two dynamical clones per computational unit, we developed a variant of the algorithm that we call the multi-clone model. The multi-clone model creates multiple dynamical clones in a single step instead of iteratively duplicating individual units, and therefore simplifies the process of keeping track of each clone’s phase shift and weight amplitude. For the multi-clone model in Figure 5, we introduced four dynamical clones (*p*1, *p*2, *p*3, *p*4) per computational unit (*p*) with paired phase shifts and balanced noise. The four clones were assigned phase shifts *δ*_*p*1_ = −*δ*_*p*2_ = *δ*_*p*3_ = −*δ*_*p*4_, where *δ*_*p*_ were drawn uniformly from [30^*°*^, 75^*°*^] for all *p*. These phase shifts were used to calculate the sinusoidal output weight profile from units in one computational unit onto units in all other computational units.

For example, the output weights from clone *p*1 onto units in other computational units is:

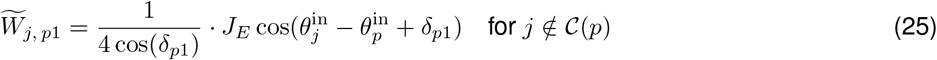

In addition, balanced noise was added to the weights such that, within each pair of clones sharing the same phase shift (e.g. *δ*_*p*1_ = *δ*_*p*3_), the noise summed to zero. For example, for clones *p*1 and *p*3, we added balanced noise such that:

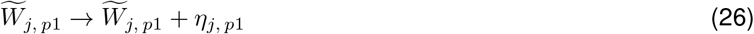

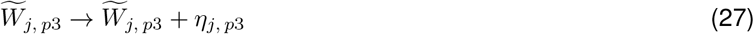

where *η*_*j, p*1_ = −*η*_*j, p*3_ is a random noise scalar drawn from 𝒩 (0, 0.2) for all shared output weights from presynaptic clones *p*1 and *p*3 onto postsynaptic unit *j*. Similarly, an independent noise term was added to the output weights from clones *p*2 and *p*4. The addition of noise to each unit’s output weights alters its underlying phase shift, which can only be recovered by averaging the weights across units that share the same phase shift. Similarly, balanced noise ensures that, when the output weights from all four clones are summed, the balanced noise cancels and the phase shifts average to zero, recovering the original symmetric connectivity, as shown in Figure 5.

#### Testing fixed points and continuous attractor dynamics

To verify that the constructed networks maintain continuous attractor dynamics, we used a two-stage procedure. First, to identify candidate fixed points along the manifold, we ran the model with a small nonzero velocity input (a “sweep”) to drive the bump around the ring and recorded the activity vector at 360 equally spaced bump angles. Second, for each candidate fixed point, we re-initialized the network at that state and simulated the network for 6 seconds with zero velocity input to confirm that the bump showed minimal drift regardless of where the bump started, as shown in Figures 4c and 5c. This is in contrast to discrete attractor networks, where the 360 candidate fixed points drift to a small number of discrete fixed points, as shown in Figure 1b.

#### Velocity input scaling

For accurate integration, the velocity input weights 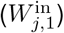 to each unit (*j*) must be scaled in proportion to each clone’s phase shift (see SI Materials Section 4.4.3). Briefly, in classical models, the asymmetric connectivity used to update the bump is parameterized as 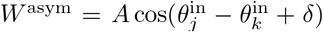, with an angular phase shift of *δ* and a recurrent weight amplitude of *A*. The network’s angular integration gain (*g*), which determines how much the bump rotates in response to a given velocity input, depends on both *δ* and *A* according to *g* = *A* sin(*δ*). Since clones with larger phase shifts have larger recurrent weights (scaled by 1/(2 cos *δ*)), their velocity input must be reduced to maintain uniform gain according to:

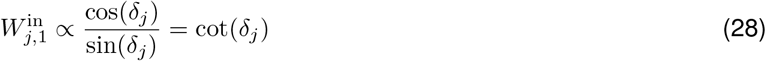

Interestingly, while this scaling relationship was derived using classical models that employ multiplicative weights, we found that applying these scaling rules to individual units in dynamical clone models results in smooth and uniform velocity integration, even though velocity inputs are additive in these networks and each unit can have a different phase shift (see SI Materials Section 4.4.3 for further explanation).

#### Initializing an unconstrained RNN model

After confirming the presence of continuous attractor dynamics, we used the resulting recurrent weight matrix to initialize an unconstrained RNN model that was used to measure the smoothness of velocity integration (in Figure 4d and Figure 5e). The output weights were initialized as cosine and sine functions of the units’ preferred directions:

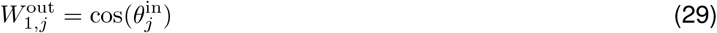

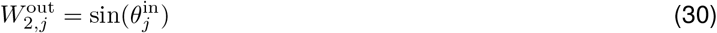

This readout extracts the first Fourier mode of the population activity vector and uses its phase as the bump position, as in classical models. The input weights that set the initial bump position, 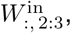, were also initialized as 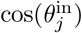 and 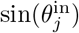 to generate a bump of population activity at the first time step. The velocity input weights were initialized according to each unit’s phase shift:

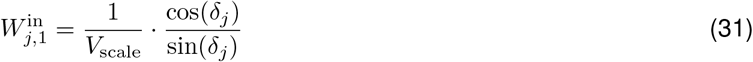

where *V*_scale_ was empirically determined by adjusting the network’s gain of integration to 1. Recurrent biases were all set to *b*_*j*_ = *c*_ff_ = 1.

### Connectome analysis (Figure 6)

To test the predictions of the dynamical clone algorithm, we analyzed the structure of recurrent connectivity between the major HD-tuned neuron types in the fly compass network.

#### Connectivity data and dynamical clone identification

Raw synaptic connectivity data were obtained from the Janelia CNS [35] and Hemibrain [33] connectomes for the major HD-tuned neuron types: Delta7 and EPG. Synapses were restricted to be within the protocerebral bridge (PB) or ellipsoid body (EB) neuropils. Neurons were assigned to computational units based on their innervation patterns in the protocerebral bridge, with angle assignments shown in Figure 6. The PB contains 18 glomeruli that can be reduced to 8 (for Delta7) or 16 (for EPG) computational units, in agreement with previous physiological data [44, 68] and our connectome-constrained RNNs (Figure 2h, middle panel). Each unit’s preferred head direction was inferred from the glomerulus it innervates, as indicated by the “instance” field returned in the neuPrint data frame. EPG and Delta7 angles were offset by 11.25 to match the measured half-glomerulus shifts [44, 68]. Neurons were then ordered within each type by increasing angle. Neurons within the same computational unit (same glomerular innervation and thus same predicted directional tuning) were considered putative dynamical clones.

#### Weight averaging to reveal hidden symmetries

We focus our analysis of hidden symmetries on Delta7-to-Delta7 connections and EPG-to-EPG connections because both of these HD-tuned populations have a high rate of putative dynamic clones whose recurrent connectivity is approximately circularly symmetric but with considerable heterogeneity. For each presynaptic unit *k* in the Delta7-to-Delta7 or EPG-to-EPG connectivity matrix (*W*), we computed the average output weight profile onto all postsynaptic units *j* grouped by their computational unit (Figure 6b and 6g, middle panels):

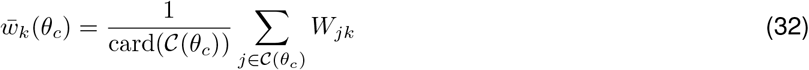

where 𝒞 (*θ*_*c*_) is the set of postsynaptic units in the computational unit with preferred direction *θ*_*c*_. Then, we shifted these profiles to align their connectivity for direct comparison (Figure 6b and 6g, right panels). For the Delta7 neurons, the symmetry of the averaged weight profile was quantified by fitting a sinusoidal function and computing the coefficient of determination (*R*^2^) of the fit. Fits were obtained by nonlinear least squares, returning per-unit amplitudes, phases, offsets, and goodness-of-fit. These fits were also performed on the average aligned weight profile across all units (Figure 6c) and on the average aligned weight profile from all units that share a computational units (black lines in Figure 6d). For EPG neurons, these phase and amplitude values were computed using a PVA whose weights came from the aligned weight profiles.

---

**Figure 2 S1:**
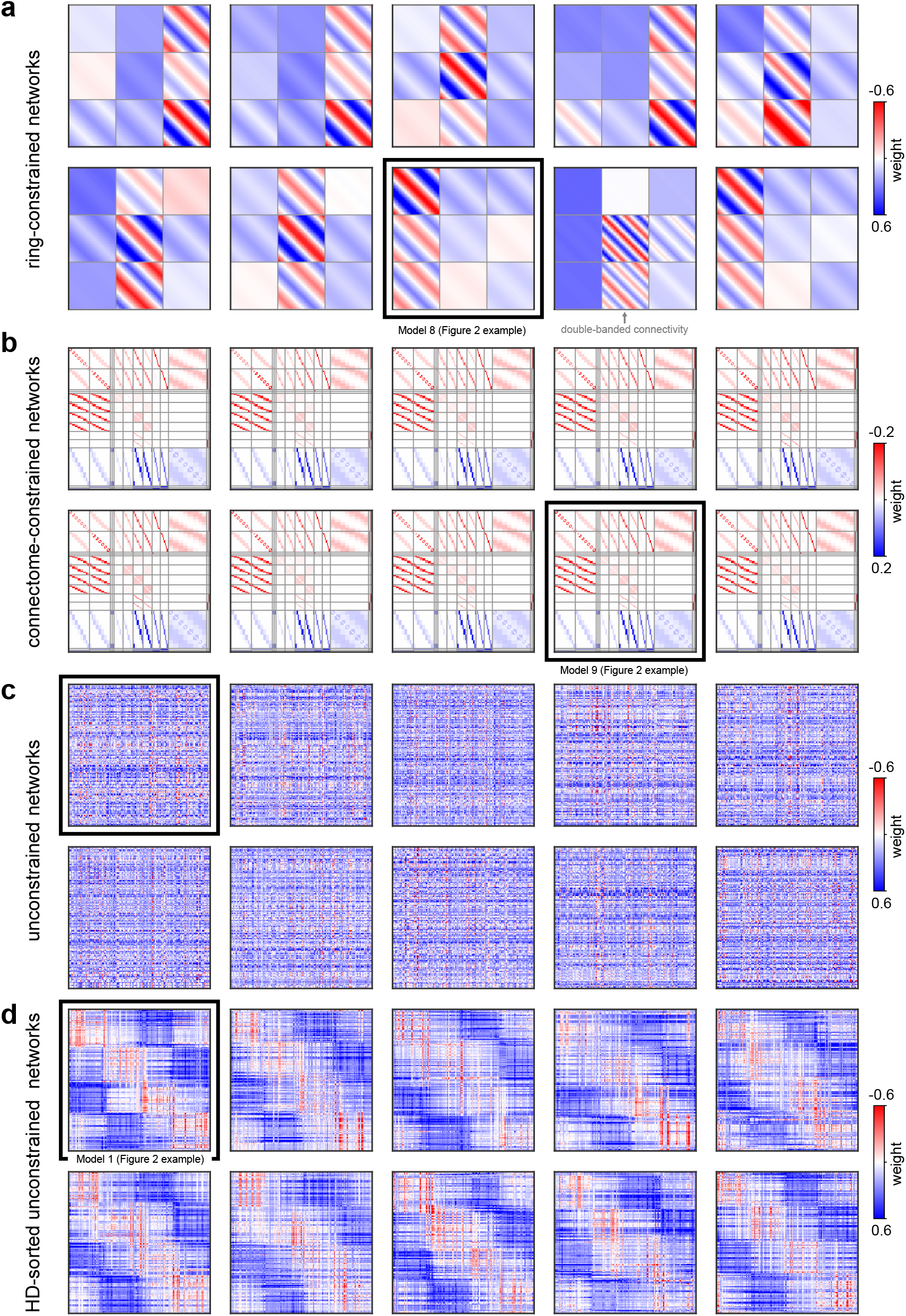
Recurrent weight matrices for all trained networks. **a)** Recurrent weight matrices for 10 ring-constrained networks. Networks are ordered 1-5 (left to right) on the top row, and 6-10 on the bottom row. The black rectangle around Model 8 marks the example model shown in the main text Fig. 2. Model 9 highlights a network that learned an approximate two-ring, two-bump solution (similar to the networks described in [12, 15, 119]). Model 4 learned an approximate two-ring, one-bump solution similar to Xie et al. [10]. The rest of the models learned approximate three-ring solutions ([8]; see Fig. S2 to S4 for additional details on these models). **b)** Same as in (a) but for connectome-constrained networks. **c)** Same as in (a) but for unconstrained networks with unsorted weights. **d)** Same as in (c) but the unconstrained weights have been sorted by the units’ preferred firing directions.

**Figure 2 S2:**
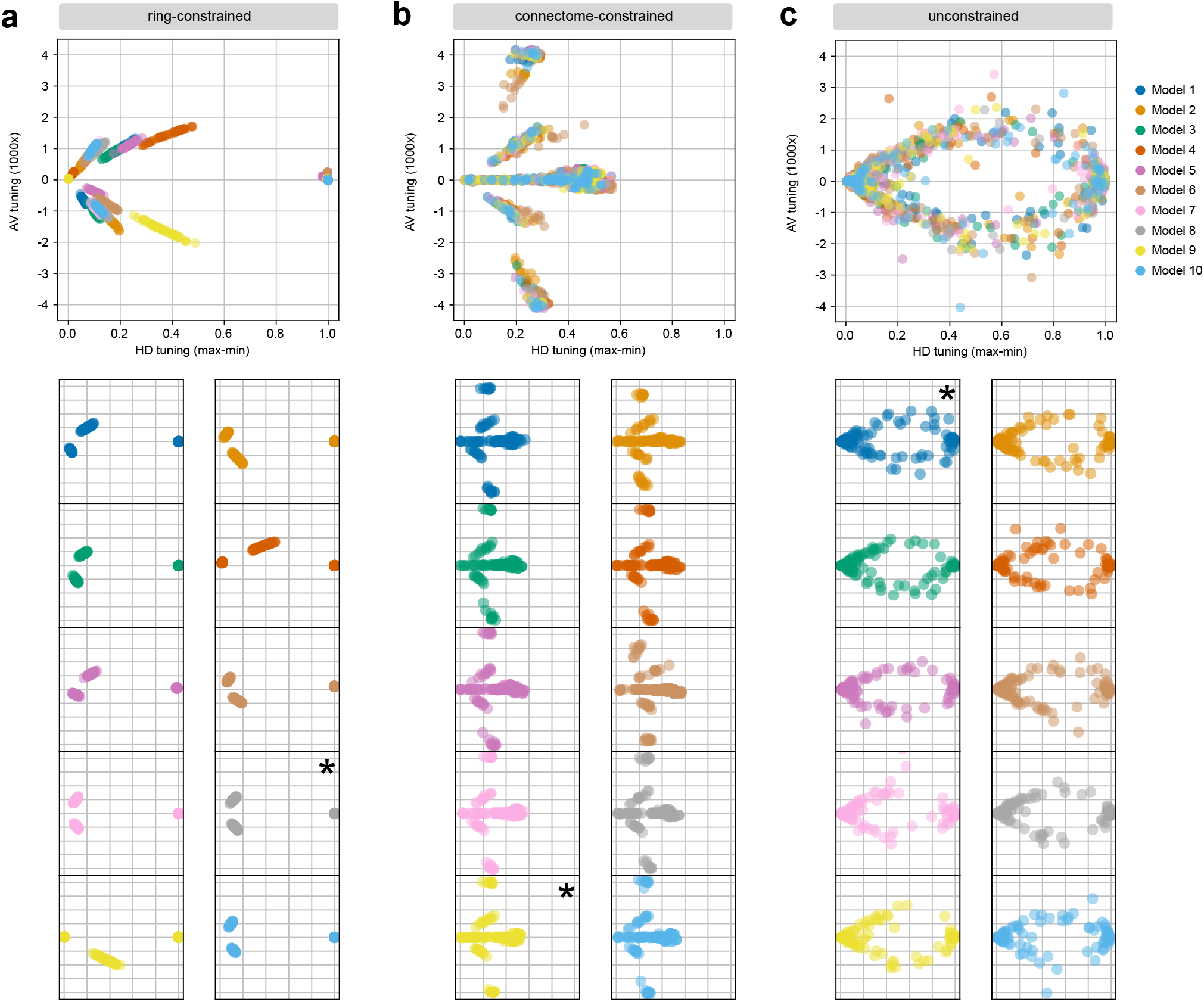
Tuning curve distributions for all trained networks. **a)** Top: tuning curve distributions for all 10 ring-constrained networks, as in Fig. 2e. Each point is a unit, and the points are colored by the network they belong to (legend shown at far right of panel c). Bottom: tuning curves distributions for the each of the 10 network shown individually instead of overlapping. Asterisks mark the networks shown in the main text Fig. 2. **b)** Same as in (a) but for connectome-constrained networks. **c)** Same as in (a) but for unconstrained networks.

**Figure 2 S3:**
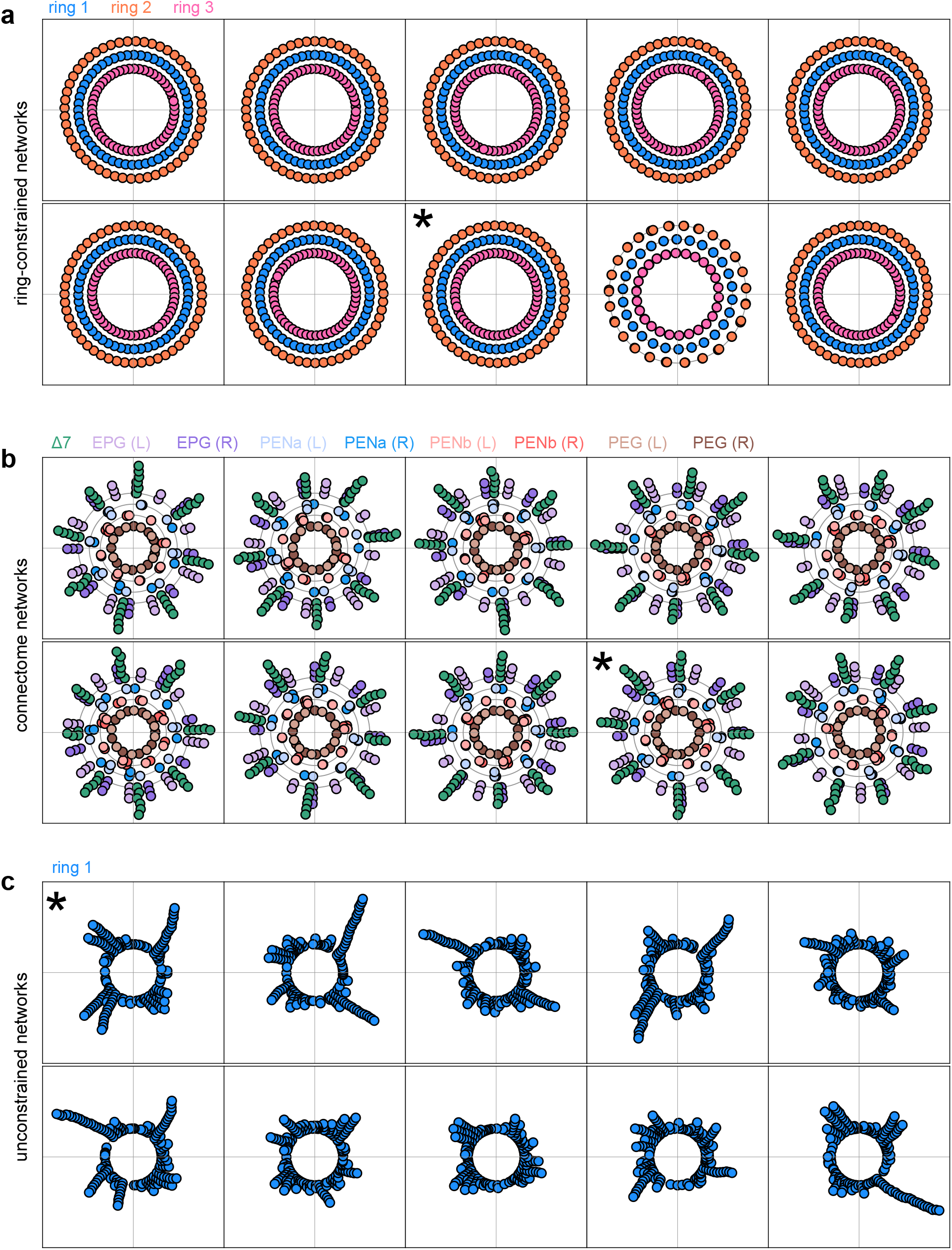
Distribution of preferred HD tunings for all trained networks. **a)** Top: distribution of preferred HD tunings across units in each of the 10 ring-constrained networks, as in Fig. 2h. Each point is a unit from single network, and points are colored by the ring they belong to. The radial distance of the points were adjusted to enable visualization of HD tunings in different rings. Networks are ordered 1-5 (left to right) on the top row, and 6-10 on the bottom row. **b)** Same as in (a) but for connectome-constrained networks. To visualize dynamical clones, units from the same cell type whose preferred HD tuning was within 5 degrees were stacked radially. **c)** Same as in (b) but for unconstrained networks. Clones whose preferred HD tuning was within 2.5 degrees were stacked radially.

**Figure 2 S4:**
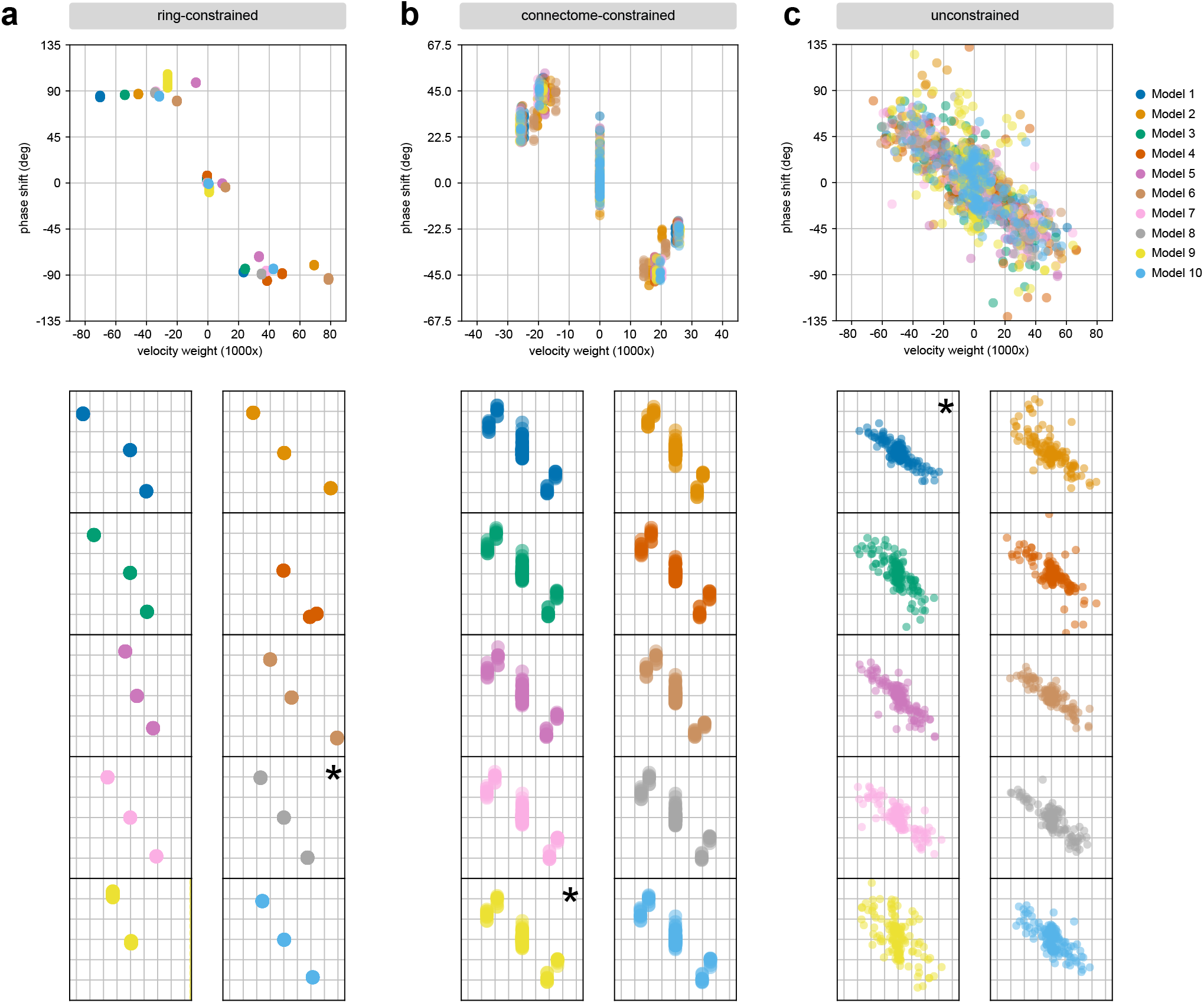
Phase shift versus velocity input strength for all trained networks. **a)** Top: phase shifts plotted as a function of velocity input strength for all 10 ring-constrained networks, as in Fig. 2i. Each point is a unit, and the points are colored by the network they belong to (legend shown at far right of panel c). Bottom: same as in the top plot but for each of the 10 networks shown individually instead of overlapping. Asterisks mark the networks shown in the main text Fig. 2. **b)** Same as in (a) but for connectome-constrained networks. **c)** Same as in (a) but for unconstrained networks.

**Figure 4 S1:**
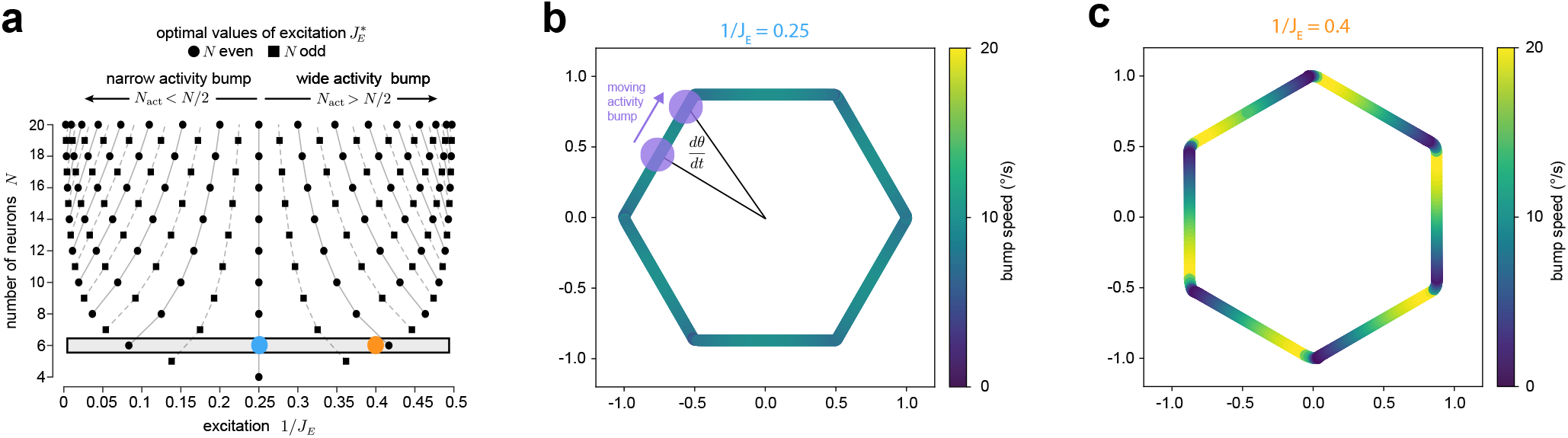
Manifold geometry affects velocity integration. **a)** Plot showing the recurrent excitation values (*J*_*E*_) that enable exact continuous ring attractor dynamics for networks of different sizes (black points), as first derived by Noorman et al. [40]. Networks whose *J*_*E*_ values are between the black points generate discrete attractors where the bump is only stable at discrete points on the ring. In the main text, we always start with the network shown by the blue dot, a continuous attractor with 1*/J*_*E*_ = 0.25 and *N* = 6 (see Methods). **b)** Because our networks use a threshold-linear activation function, the ring attractor manifold in the *N*-dimensional state space of network activity is a polygon composed of *N* linear subsystems stitched together at transitions between subsystems. For the *N* = 6 networks, this creates a hexagon embedded in a 6-dimensional state space that can be visualized by projecting activity into the 2-dimensional output space, as shown here. Specifically, the network was driven by a constant 10 degree/s velocity input, and at each point in time, the 6-dimensional firing rate vector was projected into the output space using sine and cosine basis vectors that extract the first Fourier mode of the bump. Each point is colored according to its angular velocity. Notice that points in the middle of each linear subsystem have slightly higher angular velocity than expected, and points further from the origin have slightly lower angular velocity. This is an effect of the geometry of the manifold, where points closes to the origin will change their angular position faster than points further from the origin, leading to uneven velocity integration that cannot be adjusted through changes in network weights, since it results from the choice of nonlinearity that the theory of small networks in Noorman et al. [40] relies on. Note that a similar effect of manifold geometry was shown by Darshan et al [70]. **c)** Same as in (b) but for the discrete attractor network marked by the orange point in a. In this case, as first shown by Noorman et al. [40], discrete attractor networks have highly uneven velocity integration due the presence of discrete fixed points in a bumpy energy landscape.

**Figure 6 S1:**
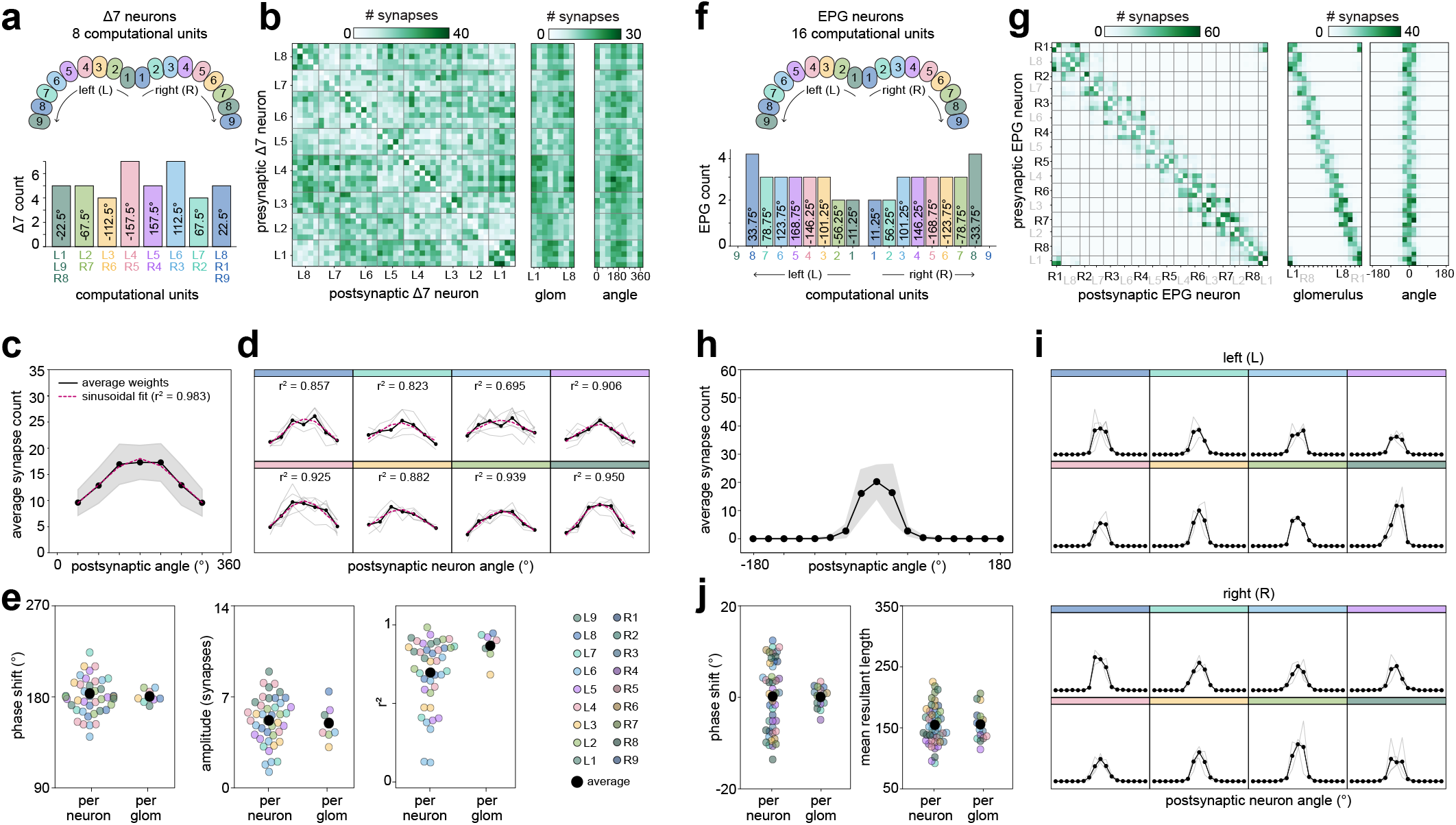
Recovering hidden symmetries in the fly connectome (Hemibrain volume). Same as Fig. 6 but for the Hemibrain volume.

**Figure 6 S2:**
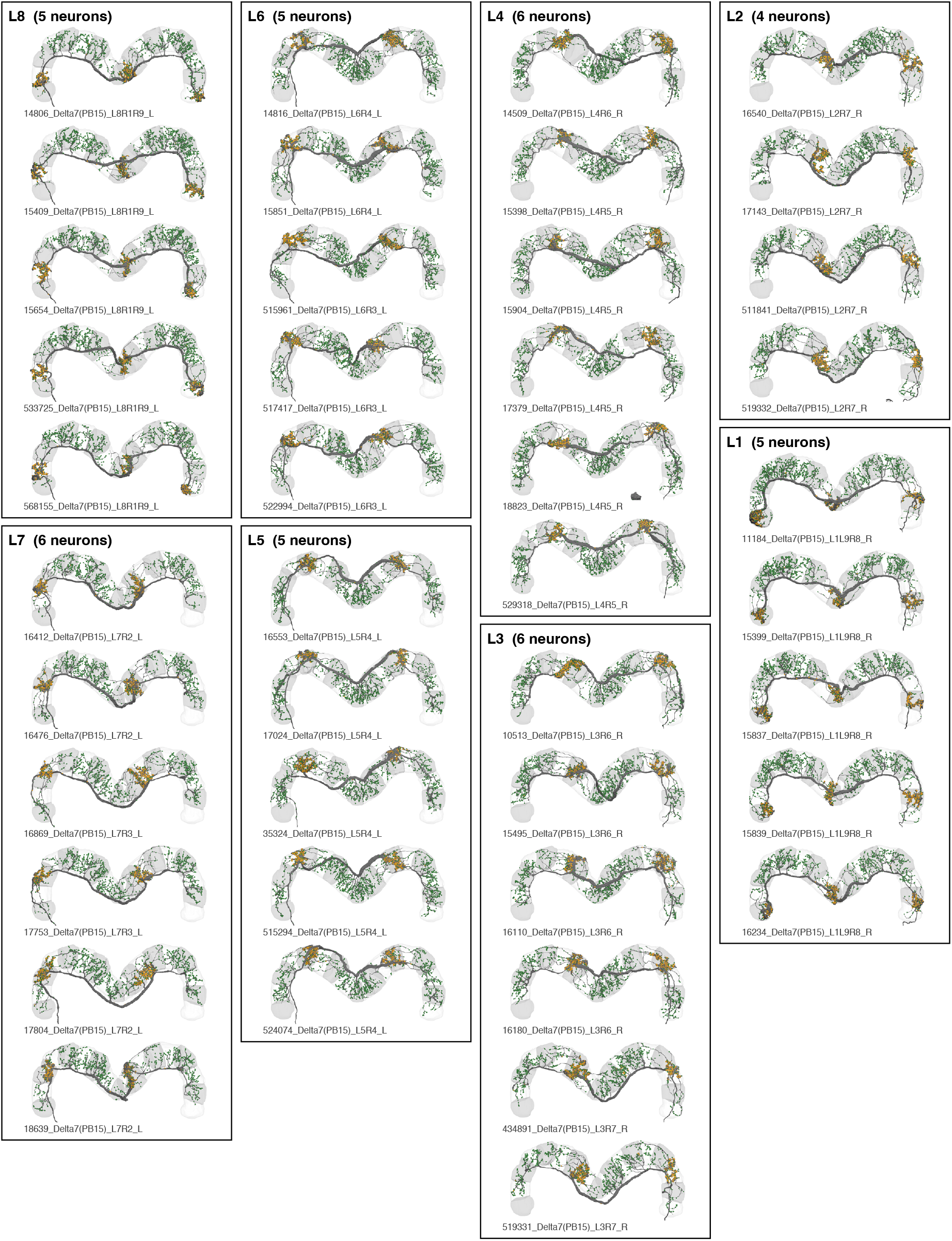
Morphological renderings of all Δ7 neurons from the CNS volume. Neurons are grouped into sets of putative dynamical clones that compose a computational unit. Postsynaptic sites that receive inputs from other Δ7 neurons are marked with green dots. Presynaptic sites that synapse onto downstream Δ7 neurons are marked with gold dots. PB glomeruli are shown in gray with alternating light/dark shading. The neurite leading to the soma has been cropped for better visualization. Each Δ7 neuron innervate 2-3 glomeruli in the PB, with neurons that innervate the same glomeruli composing a computational unit. Notice that the density of postsynaptic sites increases in an approximate sinusoidal fashion with distance from a neuron’s axonal sites. This gives rise to the Δ7 neuron’s cosine-connectivity shown in Figure 6 and Fig. 6 S1.

