## Supplemental Materials for "Hidden symmetries in network connectivity support ring attractor dynamics in the fly’s neural compass"

### From classic ring attractor models to the dynamical clone algorithm: theoretical background and derivation of key results

*Supplemental materials for:*

Hidden symmetries in network connectivity
support ring attractor dynamics in the fly's neural compass

Brad K. Hulse, P.B. Aneesh, Sandro Romani,
Vivek Jayaraman, and Ann M. Hermundstad

**5 Connection to canonical HD models** **14**

**A Reduction of firing rate equation to low modes** **15**

**B Useful integrals for the bump regime** **16**

**C Derivation of the $\theta_c$ equation** **16**

**D Clone-pair identity and gain scaling** **16**

**E Remark on exact versus approximate cloning and slope rescaling** **17**

### 1 Introduction

In this SI Materials section, we begin by providing a high-level introduction to ring-attractor networks from a historical perspective to provide theoretical background for the dynamical clones algorithm. Subsequent sections provide more detailed derivations by reviewing classic continuous ring models (Section 2) as well as more recent work that constructs continuous ring attractors using a finite number of neurons (Section 3). In the process, we highlight key theoretical results that the cloning algorithm relies on. Lastly, we provide a detailed derivation of our main theoretical contribution, the dynamical clones algorithm (Section 4).

Head direction (HD) cells are neurons whose firing rate peaks when an animal faces a particular direction; across the population, their preferred directions span the full 360 degrees of headings. HD cells were first discovered in the rodent postsubiculum [1] but have since been found in a variety of species, including primates [2], bats [3], fish [4], and insects ([5], reviewed in [6–8]). These cells maintain their directional tuning even in complete darkness, implying that the representation is generated and sustained internally by recurrent circuit dynamics rather than being driven entirely by external sensory cues (review in [6, 7, 9–11]).

Starting about five years after their discovery, a series of theoretical studies showed that ring attractor networks can elegantly account for the key properties of HD cells: a unique, persistent bump of activity whose position encodes the animal's heading, and the ability to smoothly update that position in response to angular velocity input [12–28]. Because these models were inspired by rodent cortex, where the relevant neural populations contain thousands of neurons, and because the mathematics is considerably simpler when one can replace discrete sums with integrals, the classical theory assumes an effectively infinite number of neurons arranged on a continuous ring. In this continuum limit, the firing-rate dynamics of the network evolve according to

$$\tau \frac{\partial h(\theta, t)}{\partial t} = -h(\theta, t) + \int_{-\pi}^{\pi} \frac{d\theta'}{2\pi} W(\theta - \theta') \; [h(\theta', t)]_+ + I(\theta).$$

Here, $h(\theta, t)$ is the activity (subthreshold voltage) of the neuron at angular position $\theta$, and $[\cdot]_+$ denotes the threshold-linear (ReLU) nonlinearity, $[x]_+ = \max(0, x)$, which transforms subthreshold activity into a firing rate (other nonlinearities such as sigmoids are also used in the literature and give qualitatively similar results). In firing rate models like these, each neuron's activity decays on its own (the $-h$ term), is driven by external input $I(\theta)$, and receives recurrent input from every other neuron on the ring, weighted by a perfectly symmetric connectivity kernel $W$ that depends only on the angular distance $\theta - \theta'$ between pre- and postsynaptic neurons. When the kernel has a "Mexican hat" shape, with local excitation surrounded by broad inhibition, the network spontaneously forms a localized bump of activity. Because there is an infinite number of neurons and the connectivity kernel depends only on angular differences between neurons, the bump can sit at any position around the ring with equal stability, producing the continuous ring attractor.

To update the bump's position in response to the animal's turning, a process called angular velocity integration, classical models introduce an asymmetric, or "odd," component into the connectivity [16, 23, 29]. However, this velocity-integration mechanism is typically added on top of the core attractor module rather than being intrinsic to it. In some models, separate "side rings" of neurons receive angular velocity input and project back onto the "center ring" with phase-shifted connections, as in the 3-ring model of Skaggs et al. [16]. Similarly, 2-ring models rely on balanced phase shifted connectivity between two side rings that receive left or right velocity information ([23]). In other models, the recurrent weight profile itself is modulated by velocity, as in Zhang [14], which is mathematically elegant but requires connectivity that changes in real time, a less biologically plausible mechanism. In all these formulations, velocity input strength and phase shifted connectivity are properties shared by all neurons in a given ring rather than being assigned to individual neurons.

With the discovery of HD cells in the *Drosophila* central complex [5, 30], it became clear that even small neural systems – with on the order of $\sim$100 neurons – can function as ring attractors. The fly's compass network maintains a bump of activity that tracks heading with remarkable precision, despite having far fewer neurons than classic ring attractor models assumed. This observation challenged the long-standing assumption that continuous attractor dynamics require a large number of neurons.

To bridge this gap, Noorman et al. [31] showed that it is possible to construct exact continuous attractor networks with a small number of units – as few as four – provided that the strength of recurrent excitation is tuned to specific critical values. In a finite network of $N$ neurons with evenly spaced preferred headings, the firing-rate dynamics take the same general form as the continuum equation above, but with the integral replaced by a discrete sum:

$$\tau \frac{dh_j}{dt} = -h_j + \frac{1}{N}\sum_{k=1}^{N}\Big(J_I + J_E\cos(\theta_j - \theta_k)\Big)\ [h_k]_+ + c_{\mathrm{ff}}, \qquad j = 1, \ldots, N.$$

Here, each postsynaptic neuron $j$ now sums contributions from a finite set of presynaptic inputs $k$ rather than integrating over a continuous ring. Here we've replaced the $W(\theta - \theta')$ kernel in the equation above with an explicitly cosine-shaped connectivity profile that we'll rely on throughout the sections below, where $J_E$ is the strength of recurrent excitation and $J_I$ provides broad inhibition. Regardless of the specific symmetric connectivity profile used, the finite sum generically breaks the perfect rotational symmetry of the continuum model: whereas the integral is invariant to any rotation $\psi \to \psi + \epsilon$, the discrete sum is only invariant to rotations that map the lattice of preferred headings onto itself. Put another way, in models with an infinite number of neurons, the continuous symmetry ensures that the bump shape at every angle is identical, which we refer to as a *metric ring*. In contrast, the discrete symmetry in small attractor networks ensures identical bump shapes only at $2\pi/N$ angle intervals and within such an interval, the bump shape changes, which we refer to as a *topological ring* [32]. As a result, it was thought that a naively discretized ring attractor could only support a small number of stable bump positions, rather than a true continuum, making it unclear how the fly's small attractor network could generate continuous ring attractor dynamics.

Noorman et al. [31] resolved this mystery by showing that for specific, isolated values of the excitatory coupling $J_E$, the energy landscape along the ring direction becomes exactly flat, restoring a continuous family of stable bump positions even in a network with very few neurons. As in continuous networks, velocity integration in these finite networks also relies on explicit side rings (or velocity-modulated weights), ensuring that the overall architecture is highly modular and that individual units can always be assigned to discrete rings that share the same velocity input strength and phase shift connectivity profile.

Despite their elegance, both the classical continuum models and the finite constructions of Noorman et al. [31] share an important limitation: they all require perfectly symmetric weights, and even small perturbations to these finely-tuned synaptic weights can cause the continuous ring manifold to degrade into a small number of discrete fixed points, as described in the main text. And yet, the fly compass network exhibits robust ring-attractor dynamics, despite exhibiting substantial synaptic heterogeneity that is predicted to degrade these dynamics. This is the central puzzle motivating the present work.

The key insight of this paper is that previous ring attractor models, both continuous and discrete, rely on units that uniformly tile the space of HD angles, with each angular position represented by exactly one unit. Our construction algorithm departs from this assumption by introducing *dynamical clones*: multiple units that share the same preferred heading (and therefore the same firing rate in the absence of velocity input) but can differ in their output connectivity. This redundancy overparameterizes the recurrent weight matrix, creating extra degrees of freedom that can absorb heterogeneity without disrupting the collective dynamics. The resulting networks retain *hidden symmetries*: the single-neuron weights look highly heterogeneous, but when the outputs of all clones sharing a preferred heading are summed, the original symmetric connectivity can be recovered.

A second departure from previous models is in how velocity integration is implemented. Classical architectures assign phase shifts and velocity inputs as properties of entire rings or populations. Our construction instead builds phase shifts at the level of individual neurons: each clone within a group can carry its own phase shift, and the group's phase shifts are counterbalanced so that they cancel when the clones fire at equal rates. When a velocity input differentially modulates the clones, the counterbalanced shifts produce a net odd component in the effective connectivity, driving the bump to move – exactly the mechanism needed for angular path integration. This means that velocity integration is an emergent property of the clone group rather than a feature hard-wired into a separate ring.

The remainder of this note develops these ideas in detail and is organized as follows. Section 2 reviews the canonical continuum ring model. We derive the amplitude–phase reduction that separates bump shape from bump position, and show that an odd (sine) component in the connectivity kernel directly drives the bump phase. This section establishes the mathematical language used throughout. Section 3 turns to finite networks. We explain how discretization generically breaks the continuous attractor into a small number of fixed points, and review the fine-tuned constructions of Noorman et al. [31] that restore exact continuous attractors in small networks. Section 4 introduces the clone operations that are the core contribution of the main text. We derive the conditions under which cloning preserves a continuous attractor manifold (summed-output equivalence), show that clone groups naturally accommodate balanced noise, and establish the counterbalanced phase-shift rule and the associated $1/(2\cos\delta)$ weight-scaling identity. We also derive the velocity-input scaling rule $W^{\mathrm{in}} \propto \cot\delta$ and show how multiple clones with different phase shifts reduce to an effective single-ring theory. Section 5 connects these constructions to the classical 2-ring and 3-ring HD models, showing that those architectures are special cases of the more general clone framework.

### 2 Canonical ring theory and moving bumps: a review

This section reviews the classical continuum ring attractor model [12–19]. The main results are: (i) when the network's recurrent connectivity has only zeroth- and first-order Fourier components, the full infinite-dimensional dynamics reduce to just three variables – mean activity $h_0$, bump amplitude $\rho$, and bump phase $\psi$; (ii) the even (cosine) part of the connectivity kernel controls bump shape while the odd (sine) part drives bump motion; and (iii) a phase-shifted cosine kernel provides an exact, non-perturbative way to produce the odd component needed for velocity integration. Although the notion of a bump is defined later in section 2.2 and $h_0, \rho$ and $\psi$ are defined independently of any 'bump', the bump regime will be the case of interest for us and hence we pre-emptively use this terminology. These results set up the notation and intuition needed for the finite-network and clone constructions in later sections.

#### 2.1 Canonical single ring

Canonical neural-network models of the head-direction system posit a circular ring of neurons whose connectivity mirrors the rotational symmetry of the encoded variable. In the continuum limit, the firing-rate dynamics can be written as

$$\tau \frac{\partial h(\theta,t)}{\partial t} = -h(\theta,t) + I(\theta) + \int_{-\pi}^{\pi} \frac{d\theta'}{2\pi} W(\theta,\theta')\ [h(\theta',t)]_+ \,. \tag{1}$$

Here $h(\theta,t)$ is the activity (i.e. voltage) of the neuron at angular position $\theta$, $[x]_+ = x\Theta(x)$[1] is the threshold-linear nonlinearity (which sets negative activities to zero, modeling the fact that firing rates cannot be negative), and $\tau$ is the neural time constant. The three terms on the right have a simple interpretation: the first causes activity to decay in the absence of input; the second represents external drive; and the third sums up recurrent input from every other neuron, weighted by a connectivity kernel $W$ that depends only on the angular distance $\theta - \theta'$ between presynaptic ($\theta'$) and postsynaptic ($\theta$) neurons. While the above equation is a seemingly complicated *integro-differential* equation, we will see how one can systematically solve this by going to Fourier space and cleverly choosing the form of the connectivity kernel and input.

In particular, we take the input and recurrent kernel to be restricted to the first Fourier harmonics,

$$I(\theta) = I_0 + I_1 \cos(\theta - \tilde{\psi}), \tag{2}$$

$$W(\Delta) = W_0 + W_1 \cos\Delta + C \sin\Delta, \qquad \Delta = \theta - \theta'. \tag{3}$$

The input has a uniform component $I_0$ and a directional component of strength $I_1$ centered at angle $\tilde{\psi}$, which models a sensory cue that anchors the bump. The kernel[2] has three parts: a constant $W_0 < 0$ that provides global inhibition (every neuron inhibits every other neuron by the same amount), a cosine term $W_1 > 0$ that provides structured excitation (nearby neurons excite each other more than distant ones), and an odd component $C \sin\Delta$ that will later be responsible for moving the bump. The static ring model (without velocity integration) is the special case $C = 0$. Note that since $W(\Delta)$ enters (1) via a convolution, this implies that for the choice of (3), the convolution only picks out the DC, sine, and cosine components. This simplification is key to analytically solving the dynamics in a way that that provides key conceptual insights.

In order to solve (1), it is convenient to go to Fourier space, where $h(\theta,t)$ can be expressed as a sum over harmonics (sines and cosines of different integer frequencies) with different coefficients (see appendix A). In this picture, the coefficients have the time dependence, while the harmonics themselves are independent of time. As a result, the single time derivative of $h(\theta,t)$ is replaced with the time derivatives of the infinitely many coefficients. However, since the non-decaying terms on the right hand side of (1) contain only the first harmonics from both the input (by choice from (2)) and from the convolution with the kernel, the higher harmonics of $h(\theta,t)$ necessarily decay exponentially $\sim e^{-t/\tau}$ in time. Hence, after a few timescales $\mathcal{O}(\tau)$, it is natural to look for a form of $h(\theta,t)$ that contains only the DC, sine, and cosine terms. This motivates the bump solution (ansatz),

[1] $\Theta(x)$ is the Heaviside step function, where $\Theta(x) = 1$ if $x > 0$ and $\Theta(x) = 0$ otherwise

[2] The precise signs and values of $W_0$ and $W_1$ need not be assumed *a priori*, but rather derived from imposing conditions of bump stability on their phase diagram

$$h(\theta, t) = h_0(t) + \rho(t)\cos\big(\theta - \psi(t)\big). \tag{4}$$

Here, $h_0$ is the mean activity across the ring, $\rho \geq 0$ is the bump amplitude, and $\psi$ is the bump phase (where the bump is centered on the ring manifold). This decomposition is the key simplification: instead of tracking the activity of infinitely many neurons, having a connectivity profile with just the first fourier modes, along with a DC offset, leads to a solution where the network's dynamics can be described with just three bump parameters – mean activity $h_0$, bump amplitude $\rho$, and bump phase $\psi$ – as shown below.

For the choice of the kernel (3), the integral on the right hand side of (1), can be decomposed into a sum of three integrals, each projecting $[h(\theta, t)]_+$ onto the DC, cosine and sine parts of the kernel. To project the dynamics onto these three modes, define the projections of the thresholded activity onto the DC, cosine, and sine modes:

$$g_0 = \int_{-\pi}^{\pi} \frac{d\theta'}{2\pi}\, [h(\theta', t)]_+\,, \tag{5}$$

$$g_c = \int_{-\pi}^{\pi} \frac{d\theta'}{2\pi}\, \cos\theta'\, [h(\theta', t)]_+\,, \tag{6}$$

$$g_s = \int_{-\pi}^{\pi} \frac{d\theta'}{2\pi}\, \sin\theta'\, [h(\theta', t)]_+\,, \tag{7}$$

Since $\cos\theta + i\sin\theta = e^{i\theta}$, and since exponentials are easier to work with than sines and cosines, it is simpler to use the combination $g_c + ig_s$ instead of either $g_c$ or $g_s$. Furthermore, from the ansatz (4), $h(\theta, t)$ peaks at $\psi$, and hence it will be useful to phase-shift these integrals to the bump-centered frame (multiplying by $e^{-i\psi}$). This motivates the following projection[3]:

$$g_1 = e^{-i\psi}(g_c + ig_s) = \mathfrak{Re}(g_1) + i\mathfrak{Im}(g_1). \tag{8}$$

Intuitively, $g_0$ measures the average firing rate of the network, while $g_c$ and $g_s$ (combined into $g_1$) measure the strength and orientation of the bump in the thresholded activity.

We now have all the pieces we need to work towards solving (1). For the ansatz (4), the time derivative of $h(\theta, t)$ is now replaced with time derivatives of the three parameters $h_0$, $\rho$, and $\psi$. As shown in detail in appendix A, equating the DC, cosine, and sine modes on both sides of (1) and simplifying, then gives the following equations for the $h_0$, $\rho$, and $\psi$ (in terms of the projections $g_0(h_0, \rho, \psi)$ and $g_1(h_0, \rho, \psi)$),

$$\tau\dot{h}_0 = -h_0 + I_0 + W_0 g_0 \tag{9}$$

$$\tau\dot{\rho} = -\rho + I_1\cos(\psi - \tilde{\psi}) + W_1\mathfrak{Re}(g_1) - C\mathfrak{Im}(g_1) \tag{10}$$

$$\tau\rho\dot{\psi} = I_1\sin(\tilde{\psi} - \psi) + W_1\mathfrak{Im}(g_1) + C\mathfrak{Re}(g_1). \tag{11}$$

These three equations fully describe the general form of the bump dynamics even before explicitly evaluating $g_0$ and $g_1$: the first governs the mean activity level; the second governs the bump amplitude; the third governs the bump phase. We note that these are still the relevant equations when one has a discretised version of the model, except that $g_0$ and $g_1$ are replaced by sums over discrete $\theta_i$ in place of the integral over $\theta$, as discussed in more detail in Section 3.

For the continuum model, $g_1$ is real ($g_1 = \mathfrak{Re}(g_1)$): after shifting the integration variable by $\psi$, the imaginary part is odd and integrates to zero because the thresholded bump is always left-right symmetric about its center. This is a direct consequence of the continuous rotational symmetry – one can always shift the frame of reference of $\theta$ to the bump center. Furthermore, $g_1 = g_1(h_0, \rho)$ is only a function of the amplitude, and the constant mode and the equations for the modes reduce to

$$\tau\dot{h}_0 = -h_0 + I_0 + W_0 g_0, \tag{12}$$

$$\tau\dot{\rho} = -\rho + I_1\cos(\psi - \tilde{\psi}) + W_1 g_1, \tag{13}$$

$$\tau\rho\dot{\psi} = I_1\sin(\tilde{\psi} - \psi) + C g_1. \tag{14}$$

Thus, the amplitude variables $(h_0, \rho)$ are controlled by the even part of the kernel, while the odd part directly drives the phase. This clean separation – bump shape and position are governed by different parts of the connectivity – is the central insight of the continuum theory and will remain the organizing principle throughout this note.

[3] Given a complex number $z = x + iy$, $\mathfrak{Re}(z) = x$ and $\mathfrak{Im}(z) = y$ denote the real and imaginary parts of the complex number.

#### 2.2 Bump regime and reduced equations

The ansatz (4) allows three distinct activity regimes, and it is useful to keep them separate because the nonlinearity $[\cdot]_+$ behaves differently in each case.

1. **Silent regime:** if $h_0 < -\rho$, then $h(\theta) < 0$ for all $\theta$, so $[h]_+ \equiv 0$ and the recurrent term vanishes. The network is completely silent.
2. **Fully active regime:** if $h_0 > \rho$, then $h(\theta) > 0$ for all $\theta$, so the ReLU does nothing and the network is fully active everywhere. This regime does not produce a localized bump.
3. **Bump regime:** if $|h_0| < \rho$, then only a finite arc of angles is active, and the solution is a genuinely localized bump. This is the regime of interest for HD coding.

Only the third case is of interest here. In the bump regime, since $|h_0|/\rho < 1$, we define $\theta_c \in (0, \pi)$ by

$$\cos\theta_c = -\frac{h_0}{\rho}. \tag{15}$$

The firing rate of the ansatz is then modified to be

$$[h(\theta)]_+ = \rho\,[\cos(\theta - \psi) - \cos\theta_c]_+\,, \tag{16}$$

which makes $\theta_c$ more directly interpretable than $h_0$ in the bump regime. The angle $\theta_c$ is the half-width of the active region: neurons within an angular distance $\theta_c$ of the bump center are active, and those outside this arc are silent. The thresholded activity is then nonzero only for $|\theta - \psi| < \theta_c$, and the projections can be evaluated explicitly (see appendix B) :

$$g_0 = \rho f_0(\theta_c), \tag{17}$$
$$g_1 = \rho f_1(\theta_c), \tag{18}$$

with

$$f_0(\theta_c) = \frac{\sin\theta_c - \theta_c\cos\theta_c}{\pi}, \tag{19}$$
$$f_1(\theta_c) = \frac{1}{2\pi}\left(\theta_c - \frac{\sin 2\theta_c}{2}\right). \tag{20}$$

Here, $f_0$ and $f_1$ are dimensionless functions that capture how the total output ($f_0$) and directional output ($f_1$) of the bump depend on its angular width. Both are monotonically increasing in $\theta_c$: wider bumps produce more total and directional output.

Substituting into (12)–(14) gives the reduced equations that govern bump dynamics entirely in terms of the three state variables $(h_0, \rho, \psi)$ and the single shape parameter $\theta_c$:

$$\tau\dot{h}_0 = -h_0 + I_0 + W_0\rho f_0(\theta_c), \tag{21}$$
$$\tau\dot{\rho} = -\rho + I_1\cos(\psi - \tilde{\psi}) + W_1\rho f_1(\theta_c), \tag{22}$$
$$\tau\dot{\psi} = \frac{I_1}{\rho}\sin(\tilde{\psi} - \psi) + Cf_1(\theta_c). \tag{23}$$

In this form, the dependence of the state variables on the control parameters $C$ and $I_1$ becomes clear. First, when $C = 0$ and $I_1 = 0$, the equations for $h_0$ and $\rho$ do not depend on $\psi$ at all: once a bump exists, its phase is neutrally stable, as expected for a ring attractor. Second, when $I_1 \neq 0$ but $C = 0$, the bump phase is driven toward the input phase $\tilde{\psi}$ and then stops. Third, when $C \neq 0$, the odd component of the recurrent kernel drives the phase. In the unguided case $I_1 = 0$, this gives uniform drift, while for $I_1 \neq 0$, the odd component competes with cue locking and can shift the selected phase away from $\tilde{\psi}$.

Note that in (21)-(23), we still have only 3 independent state variables, since $\theta_c$ is a function of $h_0$ and $\rho$. To keep this fact clear, one can replace the dynamical equation for $h_0$ in favour of $\dot{\theta}_c$. For the sake of completeness, we include this equation as well. Differentiating (15) and using (21)–(22) yields

$$\tau\dot{\theta}_c = \frac{I_0 + I_1\cos\theta_c\cos(\tilde{\psi} - \psi)}{\rho\sin\theta_c} + \frac{W_0 f_0(\theta_c) + W_1\cos\theta_c f_1(\theta_c)}{\sin\theta_c}. \tag{24}$$

#### 2.3 Moving bump

The key result of the previous subsection is that in the unguided case (i.e., when $I_1 = 0$) and with $C = 0$, one obtains a family of stationary bump solutions parameterized by an angle $\psi$. Adding $C \neq 0$, makes the bump move with a fixed velocity $Cf_1(\theta_c)$. Here, we motivate this fact more conceptually by showing how adding a derivative of the connectivity achieves this feat ($\sin$ being the derivative of $\cos$ is a special case). Equivalently, assume a stationary bump in the lab frame of reference (fixed by $\theta_{\rm lab}$, say). If you choose a rotating frame of reference where the bump *appears* to rotate because of relative motion (fixed by $\theta_{\rm rot}$, say), the coordinate transformation between the two frames ($\theta_{\rm lab} \to \theta_{\rm rot}$) should tell us what makes the bump move, and this also implies the presence of an additional derivative of the kernel term in the dynamics.

Suppose that for $C = 0$ and $I(\theta) = I_0$, one has a stationary bump localized at $\psi$ whose profile $h_\psi(\theta)$ solves

$$0 = -h_\psi(\theta) + I_0 + \int_{-\pi}^{\pi} \frac{d\theta'}{2\pi} W(\theta - \theta') \, [h_\psi(\theta')]_+ . \tag{25}$$

That is, at steady state, the activity is proportional to the convolution of the kernel with itself, plus a constant bias. If one now wants the same profile to move with angular velocity $\Omega$, then to first order in $\Omega$ one needs to add the transport term $-\tau\Omega\,\partial_\theta h_\psi$ on the right-hand side. Indeed, via the chain rule of differentiation,

$$\frac{d}{dt} h(\theta - \psi(t)) = -\dot\psi \, \frac{\partial}{\partial\theta} h(\theta - \psi(t)). \tag{26}$$

In other words, a bump that is rigidly translating around the ring at speed $\dot\psi$ experiences an effective drive proportional to the spatial derivative of its own profile. Thus, a term proportional to $-\partial_\theta h_\psi$ directly drives the stationary bump. Equivalently, one may move $\tau\Omega\,\partial_\theta h_\psi$ to the left-hand side and interpret it as a co-moving derivative (with the coordinate transformation $\theta_{rot} = \theta_{lab} + \Omega t$). This spatial derivative of its own profile is equivalent to the convolution with the *derivative* of the stationary kernel with the activity from (25), and this is what needs to be added to the original kernel to have a moving bump.

For a general kernel, shifting the kernel by a small amount produces precisely such a term:

$$W(\Delta - \delta) = W(\Delta) - \delta\,\partial_\Delta W(\Delta) + O(\delta^2), \tag{27}$$

where we perform a Taylor series expansion in $\delta$ and retain only the leading order term. So, to leading order, adding the derivative of the kernel to itself is equivalent to a small phase shift in the recurrent interactions. The cosine kernel is especially useful because this statement can be made exact, as we show next.

#### 2.4 Exact phase-shift identity for the cosine kernel

For a cosine connectivity kernel, shifting the connectivity by an angle $\delta$, no matter how large, only introduces an additional sine term. The higher order terms effectively modulate the coefficients of the cosine and sine terms, thereby maintaining the exact functional form of the kernel as a sum of cosine and sine for any phase-shift $\delta$. For

$$W_\delta(\Delta) = W_0 + W_1 \cos(\Delta - \delta), \tag{28}$$

we have the exact decomposition

$$W_\delta(\Delta) = W_0 + J\cos\Delta + C\sin\Delta, \qquad J = W_1\cos\delta, \qquad C = W_1\sin\delta = J\tan\delta. \tag{29}$$

This is the non-perturbative version of (27) – all the higher order terms in $\delta$ of the Taylor expansion can be explicitly resummed. The reduced equations are therefore still (21)–(23), but with the even coefficient $W_1$ replaced by $J$ and the odd coefficient given by $C = J\tan\delta$. Intuitively, a phase shift $\delta$ "rotates" the cosine kernel, and the projection of this rotated kernel onto $\cos\Delta$ and $\sin\Delta$ gives even and odd components whose ratio is $\tan\delta$.

In the unguided case $I_1 = 0$, the bump velocity is

$$\tau\dot\psi = J\tan\delta\, f_1(\theta_c). \tag{30}$$

On the ring manifold, when $\dot\rho = 0$, $f_1(\theta_c) = 1/J$ (from (22)) and hence implies $\tau\dot\psi = \tan\delta$, as it should. This clarifies the standard intuition: larger phase shifts increase the odd-to-even ratio, but they also reduce the even part $J = W_1\cos\delta$, unless one compensates by increasing the overall coupling strength. Note that choosing $\delta = 90^\circ$ implies no recurrent excitation and only a rotational drive, which destroys the ring manifold. This is why phase shifts are kept below $90^\circ$ in the main text examples. Similarly, in the main text, we want to move the bump with unit speed, and hence scaling the velocity input by $\cot\delta$ can help achieve this, as described next.

##### 2.5 Exact scaling of a copy

Because the threshold-linear nonlinearity $[\cdot]_+$ is positively homogeneous (i.e., $[cx]_+ = c[x]_+$ for $c > 0$), multiplying all of a neuron's activities by a constant positive factor $v$ produces another valid solution with rescaled drive. This property will be essential for the clone construction, where it formalizes how a clone's output can be uniformly scaled up or down.

Because the ReLU is positively homogeneous, any constant positive multiple of a solution is again a solution with rescaled drive. Concretely, suppose $h_\psi^{(0)}(\theta, t)$ solves

$$\tau \frac{\partial h_\psi^{(0)}(\theta, t)}{\partial t} = -h_\psi^{(0)}(\theta, t) + I_0 + \int_{-\pi}^{\pi} \frac{d\theta'}{2\pi} W(\theta - \theta') \left[ h_\psi^{(0)}(\theta', t) \right]_+ . \tag{31}$$

If $v > 0$, then

$$h_\psi^v(\theta, t) = v h_\psi^{(0)}(\theta, t) \tag{32}$$

solves the same equation, with $I_0$ replaced by $vI_0$. Equivalently, for $I_0 \neq 0$ and $I_0 + \alpha$ of the same sign as $I_0$,

$$h_\psi^\alpha(\theta, t) = \left(1 + \frac{\alpha}{I_0}\right) h_\psi^{(0)}(\theta, t) \tag{33}$$

is exact. This means that an additive change $\alpha$ to the external drive produces a *multiplicative* change in the entire activity profile. This exact scaling statement is useful because later on, we will need a multiplicative velocity modulation via an additive velocity addition – the clone construction exploits this correspondence to convert differential velocity inputs into differential gain modulation of clone pairs.

In summary, in Section 2, we've show that the continuum ring attractor separates cleanly into amplitude (shape) and phase (position) dynamics (where the even part of the connectivity controls amplitude and the odd part drives phase velocity), a phase-shifted cosine kernel provides the odd component exactly, and the ReLU nonlinearity allows additive drive changes to act as multiplicative gain changes. These are the building blocks we will need for the dynamical clone algorithm. We turn to that construction next, but first, we need to discuss how the continuum theory extends to finite networks.

#### 3 Small discrete networks and computational units: metric vs. topological rings

This section addresses what happens when we move from the idealized continuum model to a network with a finite number of neurons, as is relevant for the fly compass system. The main takeaways are: (i) a naive discretization generically breaks the continuous attractor into a small number of stable fixed points; (ii) despite this, specially-tuned finite networks can support exact continuous attractors, as shown by Noorman et al. [31]; and (iii) in these finite networks, the bump shape may vary slightly as the bump moves around the ring, giving a *topological* rather than *metric* ring.

##### 3.1 Generic discretized rings

While the continuum model is analytically convenient, the main text starts from discrete threshold-linear networks that match the small size of the fly's ring attractor network. If one discretizes the continuum ring on lattice points $\theta_i = 2\pi i/N$ and chooses

$$W_{ij} = \frac{1}{N}\Big(W_0 + W_1 \cos(\theta_i - \theta_j)\Big), \tag{34}$$

then the exact $SO(2)$ symmetry of the continuum theory is reduced to the finite cyclic symmetry $C_N \cong \mathbb{Z}_N$. Intuitively, the continuum ring has no preferred lattice positions; the bump can sit anywhere. But a finite ring of $N$ neurons has $N$ preferred lattice positions, just as a wheel with $N$ teeth tends to rest with one tooth at the bottom. Furthermore, a lattice in position space imposes a frequency cutoff on the higher modes, but these higher frequency modes had anyway decayed, even in the continuum model. Therefore, the modes $h_0, \rho$, and $\psi$ are still the relevant modes even in the discrete case, and these modes take on continuous values, even though we have a finite number of neurons. The ansatz is then modified to be:

$$H_j = h(\theta_j) = h_0 + \rho \cos(\theta_j - \psi), \tag{35}$$

where we capitalize the left-hand side (position space) so as not to be confused with the frequency space[4]. The equations satisfied by the relevant modes are still the same as (9)–(11) but with the integrals over $\theta$ replaced by sums over $i$:

$$g_0 = \frac{1}{N}\sum_{j=1}^{N}[H_j]_+\,, \qquad g_c = \frac{1}{N}\sum_{j=1}^{N}\cos\theta_j\;[H_j]_+\,, \qquad g_s = \frac{1}{N}\sum_{j=1}^{N}\sin\theta_j\;[H_j]_+\,. \tag{36}$$

This change from $SO(2)$ to $\mathbb{Z}_N$ (continuum to discrete) has important implications for the dynamics of the network. In the continuum, the quantity $g_1$ defined in (8) is real for every phase $\psi$, and therefore $\dot{\psi} = 0$ for every $\psi$ in the absence of an external input. Furthermore, the phase decouples cleanly from the bump shape. In the discrete network, the analogous finite sum is generally not purely real for arbitrary $\psi$; its imaginary part vanishes automatically only at lattice-aligned phases, and the real part is independent of $\psi$ only when the lattice and activity profile are aligned as well. As a result, $\dot{\psi} = 0$ only at $N$ stable and $N$ unstable phases. Intuitively, when the bump is centered on a neuron's ring location (i.e., a lattice-aligned phase), the forces from the left and right neighbors balance by symmetry, and the bump stays put. When the bump is centered between neurons, the forces are generically unbalanced, and the bump drifts toward the nearest lattice position. Therefore, if one insists on a rigidly translated bump profile, the finite network generically prefers a discrete set of phases rather than a true continuum.

Could there be a way to overcome this apparent limitation of the discrete networks? Since the sums (i.e., the analogue of equations (9)–(11)) also depend on bump shape parameters ($\rho$ and $\theta_c$), perhaps $\dot{\psi}$ could equal $0$ for all $\psi$, in a manner that depends on bump shape. If so, then once one allows the bump shape to change slightly with position around the cycle, a finite network can still support a connected one-dimensional family of fixed points. This is no longer a metric ring in the strict continuum sense, because the state at phase $\psi$ is not obtained by rigidly translating a single profile. But it is still a *topological* ring: a closed, one-dimensional family of states carrying the same circular variable.

#### 3.2 Fine-tuned exact rings in small networks

As alluded to in the section above, finite network size by itself does not destroy continuous attractors. Instead, small networks can be tuned to support an exact one-dimensional continuous family of fixed points, as was shown recently for threshold-linear networks by Noorman et al [31]. These constructions are the natural finite starting point for the clone operations of the main text.

More concretely, for an $N$-neuron ring with evenly spaced preferred headings and symmetric connectivity of the form

$$W^{\text{sym}}_{jk} = \frac{1}{N}\Big(J_I + J_E\cos(\theta_j - \theta_k)\Big), \tag{37}$$

Noorman et al. [31] showed that the curvature of the energy function along bump orientation vanishes only at $N-3$ isolated values of the tuned excitation $J_E$, indexed by the number of active neurons $N_{\text{act}} \in [2, N-2]$. One convenient form of the result is

$$\frac{1}{J^{*}_{E,N_{\text{act}}}} = \frac{1}{4} + \frac{1}{2N}\left(\tilde{n} + \frac{\sin(2\pi\tilde{n}/N)}{\sin(2\pi/N)}\right), \qquad \tilde{n} = N_{\text{act}} - \frac{N}{2}. \tag{38}$$

This shows that for a network of size $N$, there are $N-3$ special values of the excitatory coupling strength $J^{*}_{E}$, each corresponding to a different number of simultaneously active neurons $N_{\text{act}}$. At each of these special values, the energy landscape along the ring direction is exactly flat, meaning the bump can sit at any orientation with equal stability – even in a network with as few as four neurons. Away from these special values, the landscape is bumpy and the networks revert to discrete attractors with a small number of preferred bump orientations.

These are the fine-tuned starting networks shown schematically in Figure 4 S1a: for each allowed active-set size, there is one isolated excitation value that makes the ring manifold exact. The clone operations of Section 4 should therefore be read as an equivalence statement: if the parent network is tuned to one of these exact manifolds, then summed-output equivalence preserves that same manifold in the cloned network because every clone group, whose members share the same firing rate in the absence of input, reproduces the parent's total recurrent drive. The clone operations do not, by themselves, create an exact continuous attractor from a generic non-optimal parent.

What is lost generically in finite networks is not continuity per se, but the simple continuum symmetry of rigid translation. A specially tuned finite network can still have an exact neutral direction, even though the state is not

[4] Noorman et al. notation is the opposite, since they purely work in the discrete setup without comparing to the continuum

obtained by translating a single fixed profile. This generic lattice-pinning statement should not be confused with the fine-tuned small-network constructions above. The latter are exact finite continuous attractors; the former are what one expects from a naive discretization of the continuum kernel.

The same point clarifies the role of standard pinning inputs of the form

$$I_i = I_0 + I_1 \cos(\theta_i - \tilde{\psi}). \tag{39}$$

In a perfectly symmetric continuum ring, such an input can pin the bump at any desired phase. In a generic finite ring, the same input instead competes with the lattice preference. Unless the network is specially tuned, the selected state is typically the nearest available phase, together with a small shape deformation.

In summary, in Section 3 we have shown that moving from infinitely many neurons to finitely many generically breaks the continuous attractor into a discrete one. But this breakage is not inevitable – Noorman et al. [31] showed that, for isolated values of the excitatory coupling, even very small networks can support exact continuous attractors. These fine-tuned finite networks are our starting point. The challenge addressed in the next section is how to introduce redundancy and heterogeneity into these networks without destroying their continuous attractor property.

### 4 The dynamical clone algorithm

This section presents our core theoretical contribution: the clone construction that enables heterogeneous connectivity in continuous attractor networks. We first establish the general algebraic conditions (summed-output equivalence) under which cloning a neuron preserves the network's fixed points and dynamics. We then apply this construction to the fine-tuned finite ring attractors from the previous section, showing how counterbalanced phase shifts with the $1/(2\cos\delta)$ scaling rule allow clone pairs to exactly reproduce the parent unit's structured output while also enabling velocity integration through differential gain modulation. We also show how balanced noise can be absorbed by clone groups, and how the effective dynamics of a multi-clone network reduce to the same single-ring theory.

| Symbol | Meaning |
|---|---|
| $h_i$ | voltage of unit $i$ in the original network, with *lowercase latin* subscripts |
| $h_A$ | voltage of unit $A$ in the cloned network, with *uppercase latin* subscripts |
| $W_{ij}$ | Connectivity of original network, $i \leftarrow j$ |
| $W_{AB}$ | Connectivity of m-plicated network, $A \leftarrow B$ |
| $p\mu$ | *Greek* subscript indexes the cloned neurons of neuron $p$ from the original network |

**Table 1:** Summary of notation used in the text.

#### 4.1 Setup

We now leave the continuum picture and work directly with finite networks. Consider a rate network of $N$ neurons with state $h \in \mathbb{R}^N$, nonlinearity $\phi : \mathbb{R} \to \mathbb{R}$ applied elementwise, and dynamics

$$\tau \dot{h}_i = -h_i + \sum_j W_{ij}\, \phi(h_j) + b_i. \tag{40}$$

Here, the first term causes each neuron's activity (voltage) to decay toward zero in the absence of input, the second sums recurrent contributions from every other neuron $j$, weighted by $W_{ij}$, and the third adds a constant external bias $b_i$. Assuming there exist steady states, they solve

$$h_i = \sum_j W_{ij}\, \phi(h_j) + b_i. \tag{41}$$

From these starting networks, we construct a larger network by *m-plicating* a neuron $i$ (cloning into $m_i$ copies) and redistributing weights in a way that preserves the original fixed points under a natural embedding map (from

$\mathbb{R}^N \to \mathbb{R}^{m_1+m_2+\cdots+m_N}$). The idea is to create redundancy: the new network has more neurons than the original, but if the clones are initialized identically and their weights satisfy certain sum rules, then every clone has the same activity as its parent unit in the absence of input, and their summed output onto every postsynaptic neuron is the same as the output from the original unit, before it was cloned. This produces networks with no manifest symmetry in the recurrent weight (e.g., no obvious circulant structure) that are nevertheless *fixed-point equivalent* to symmetric networks. By this, we mean that the cloned network is constructed so that the fixed-point structure of the ring manifold is the same as in the original (uncloned) symmetric network; each fixed point of the original network maps to a fixed point of the cloned network in which all clones of a given parent unit share that parent's firing rate. The new network has neurons $> N$ with the following dynamics:

$$\tau\dot{\widetilde{h}}_A = -\widetilde{h}_A + \sum_B \widetilde{W}_{AB}\,\phi(\widetilde{h}_B) + \tilde{b}_A. \tag{42}$$

Here, the tilde notation distinguishes the cloned network from the original: $\widetilde{h}_A$ denotes the activity of unit $A$ in the cloned network, $\widetilde{W}_{AB}$ is the recurrent weight from presynaptic clone $B$ to postsynaptic clone $A$ in the cloned network, and $\tilde{b}_A$ is the bias for unit $A$ in the cloned network. The capitalized indices $A$ and $B$ range over all units in the cloned network, in contrast to lowercase $i$ and $j$, which range over units of the original network. The steady states are given by:

$$\widetilde{h}_A = \sum_B \widetilde{W}_{AB}\,\phi(\widetilde{h}_B) + \tilde{b}_A. \tag{43}$$

#### 4.2 Motivating example – 1 neuron duplication

As a motivating example (main text Figure 4), consider duplicating a single neuron (say neuron 1, without loss of generality) in a ring attractor network. The old network has $N$ neurons while the new network has $N+1$ neurons. All the unduplicated neurons have identical activity to the old network if the output from the cloned neurons, $a$ and $b$, sum to the output from neuron $1$. And for the duplicated neurons themselves, if we allow the input to $a$ and $b$ to be identical to the input to $1$, then their firing rates are also identical to the old network. Note that this means the self coupling from $1$ to itself in the old network can be decomposed into the self coupling for $a$, as $a \to a$ and $b \to a$ (likewise for $b$). This duplication idea can be generalized readily.

#### 4.3 Cloning and m-plication

More generally, consider a neuron $p$, and consider cloning it into $m$ copies, indexed by the set $\mathcal{C}(p) = \{p\mu\} = \{p1, p2, \ldots, pm\}$. That is, we reserve *latin* $i, j$ for the unduplicated neurons and suffix *greek* $\mu, \nu$ to $i, j$ to indicate duplicated neurons. Consider further replacing just 1 of the neurons (neuron $p$, say) of the original network with the cloned copies whose activity are forced to be the same ($h_{p\mu} = h_i\ \forall \mu \in \mathcal{C}(p)$). This implies the following set of necessary and sufficient conditions:

(C1) **There must be no change among non-cloned neurons:**

$$\widetilde{W}_{ij} = W_{ij} \quad \text{for all } i, j \neq p, \quad \text{and } \widetilde{b}_i = b_i \quad \text{for all } i \neq p. \tag{44}$$

(C2) **Each clone must receive identical inputs:**

$$\widetilde{W}_{p\mu,\,j} = W_{pj} \quad \forall \mu \in \mathcal{C}(p). \tag{45}$$

This means that every clone receives the same input from neurons outside its clone group that the parent neuron received.

(C3) **Outputs from the original neuron must be split among the m-plicates:** for each unduplicated postsynaptic neuron $i \neq p$,

$$\sum_{\mu \in \mathcal{C}(p)} \widetilde{W}_{i,\,p\mu} = W_{ip}. \tag{46}$$

This is the summed-output rule: the total output from all clones onto any given postsynaptic neuron must equal the output from the original parent.

(C4) **Duplicated block must preserve self-interactions:** for each $\mu$,

$$\sum_{\nu \in \mathcal{C}(p)} \widetilde{W}_{p\mu,\,p\nu} = W_{pp}, \tag{47}$$

and $\widetilde{b}_{p\mu} = b_p$ for all $\mu$. The total coupling from the clone group onto each member of the group must match the original self-coupling.

If the clones are initialized to have the same activity as the parent clones, then assuming (C1)–(C4), each clone in a group satisfies the same differential equation, and therefore

$$\tilde{h}_{p\mu}(t) = h_p(t) \quad \text{for all } \mu \in \mathcal{C}(p). \tag{48}$$

This is because every postsynaptic clone in a group receives exactly the same recurrent drive from the network as was received by its uncloned parent unit. Although the above conditions were described for a 1-step m-plication (that is, m-plicating a single neuron), one could have chosen a different neuron ($q$, say) and shown it was equivalent to the uncloned network under the same conditions for $q$. Transitively, multiple neurons could be simultaneously cloned, and this construction always holds. This generality is what enables the dynamical clone algorithm to be applied to any recurrent weight matrix. The conditions (C1) – (C4) can be reduced to two simple equations: $\widetilde{b}_{i\mu} = b_i$ and

$$\sum_{\nu \in \mathcal{C}(j)} \widetilde{W}_{p\mu,\,j\nu} = W_{pj} \quad \forall \mu \in \mathcal{C}(p). \tag{49}$$

We get (C4) when $j = p$, (C3) when $j \neq p$ and $\mathcal{C}(p) = \{p\}$, (C2) when $\mathcal{C}(j) = \{j\}$, and (C1) when $\mathcal{C}(p) = \{p\}$, $\mathcal{C}(j) = \{j\}$. Equation (49) states that for any clone $p\mu$ of neuron $p$, the sum of weights it receives from all clones of any neuron $j$ must equal the original weight $W_{pj}$. This single equation encapsulates all four conditions above.

Note that, if the outgoing weights are divided equally among the $m$ clones, each carries $\widetilde{W}_{i,\,j\mu} = \dfrac{W_{ij}}{m}$. Furthermore, notice that only the *sum* of within-group couplings is fixed by (49). Individual self-connections may therefore be weak or even absent, so long as the total within-group coupling reproduces the parent's self-coupling.

Moreover, if the original network admits fixed points $h_i^*$, then $\tilde{h}_{i\mu}^* = h_i^*$ are also fixed points of the network that contains dynamical clones. This means that if the original network had a continuous attractor manifold, the cloned network also has the same manifold as a continuous attractor, provided that the new manifold directions introduced by the cloning procedure are also stable.

#### 4.4 Dynamic clones in finite size ring attractors

We now apply the dynamical clone algorithm to the fine-tuned finite ring attractors. The key result is that counterbalanced phase shifts, when combined with the $1/(2\cos\delta)$ scaling rule and appropriate velocity inputs, allow clone pairs to preserve the continuous attractor while enabling angular velocity integration.

Using the construction defined above, we clone the fine-tuned networks of Sec. 3.2. Because these networks admit a continuous attractor manifold that is topologically a ring, these new cloned networks have the same attractor manifold. These are the networks considered in the main text. That is, we start with connectivity of the form,

$$W_{jk}^{\text{sym}} = \frac{1}{N}\Big(J_I + J_E \cos(\theta_j - \theta_k)\Big), \tag{50}$$

and perform the cloning operations. Note that, while one can include the unstructured inhibition ($J_I$) when cloning, it is more interesting to focus on the structured directional component, i.e., the cosine part alone. The reason is that the cosine component is what carries the directional information and determines the bump's position, while the uniform inhibitory component $J_I$ simply sets the overall excitability and can be split equally among clones without any special scaling.

##### 4.4.1 Counterbalanced phase shifts and the $1/(2\cos\delta)$ rule

In this section, we show that two clones with equal-and-opposite phase shifts $\pm\delta$, whose output weights are each scaled by $1/(2\cos\delta)$, exactly reproduce the parent's structured output when their firing rates are equal. When a velocity input differentially modulates the two clones, the same pair generates an odd (sine) component that drives bump motion.

While the results of the previous sections suggest that we can allow for arbitrary sloppiness in the output weights of the clones so long as their sum is kept fixed, here we focus on *phase-shifts* with a view towards angular velocity integration. The main-text phase-shift operation acts on the structured directional component of a unit's output profile. We write that parent profile as

$$W_{\text{str}}^{(0)}(\Delta) = J_E \cos\Delta. \tag{51}$$

Any common, direction-independent term is handled separately and does not enter the algebra below. If we replace the parent unit by two clones that have opposite phase shifts $\pm\delta$ and structured output profiles given by:

$$W_{\text{str}}^{(\pm)}(\Delta) = \frac{J_E}{2\cos\delta}\cos(\Delta \mp \delta), \tag{52}$$

then

$$W_{\text{str}}^{(+)}(\Delta) + W_{\text{str}}^{(-)}(\Delta) = J_E \cos\Delta, \tag{53}$$

because $\cos(\Delta-\delta)+\cos(\Delta+\delta) = 2\cos\Delta\cos\delta$. This is the algebraic content of the $1/(2\cos\delta)$ rule used in the main text: equal and opposite phase shifts can be inserted without changing the summed structured output, provided the shifted profiles are scaled by $1/(2\cos\delta)$. The factor of $1/2$ accounts for splitting one neuron into two, and the factor of $1/\cos\delta$ compensates for the fact that shifting a cosine reduces its projection onto the original unshifted cosine.

If the two clones are then differentially modulated with a relative imbalance $m$, so that the shifted profiles carry gains $1+m$ and $1-m$, the effective structured output becomes

$$\begin{aligned} W_{\text{str}}^{\text{eff}}(\Delta) &= \frac{J_E(1+m)}{2\cos\delta}\cos(\Delta-\delta) + \frac{J_E(1-m)}{2\cos\delta}\cos(\Delta+\delta) \\ &= J_E\cos\Delta + mJ_E\tan\delta\,\sin\Delta. \end{aligned} \tag{54}$$

Thus, the equal-gain pair exactly reproduces the parent structured output, while a differential gain converts the same pair into an effective odd first harmonic with coefficient

$$C_{\text{eff}} = mJ_E\tan\delta. \tag{55}$$

This is precisely the term that drives bump motion in the reduced phase equation (23). The imbalance $m$ is proportional to the angular velocity input, so the bump velocity is proportional to the angular velocity – exactly what is needed for path integration.

A common inhibitory or background term can simply be copied or split equally across clones without the $1/(2\cos\delta)$ factor, since the scaling rule above is only needed to preserve the directional first harmonic.

##### 4.4.2 Many clones and effective single ring reduction

The construction described above extends seamlessly from a single clone pair to an arbitrary number of shifted populations. Regardless of how many clones or auxiliary populations contribute, the effective connectivity always decomposes into even and odd first harmonics, so the full multi-clone network reduces to the same single-ring theory derived in Section 2.

Nothing essential changes for more populations. If a clone or auxiliary population $a$ contributes a structured first harmonic $A_a\cos(\Delta-\delta_a)$ with gain $v_a$, then

$$W_{\text{str}}^{\text{eff}}(\Delta) = \sum_{a=1}^{n} v_a A_a \cos(\Delta-\delta_a) = J^{\text{eff}}\cos\Delta + C^{\text{eff}}\sin\Delta, \tag{56}$$

with

$$J^{\text{eff}} = \sum_{a=1}^{n} v_a A_a\cos\delta_a, \qquad C^{\text{eff}} = \sum_{a=1}^{n} v_a A_a \sin\delta_a. \tag{57}$$

Any common inhibitory or background term is added separately. Thus, apparently different multi-ring, clone-group, or heterogeneous-unit architectures collapse to the same first-harmonic reduced theory: the even part controls bump shape and the odd part controls phase velocity. This universality has an important implication: it means that no matter how complex the clone structure, the network's attractor dynamics can always be understood through just two numbers, $J^{\text{eff}}$ and $C^{\text{eff}}$.

##### 4.4.3 Angular velocity integration and velocity input scaling

Our networks provide angular velocity input to each unit as an additive term with a per-unit scaling that depends on each unit's phase shift. Clones with different phase shifts must receive different velocity input weights to ensure that all clone pairs contribute equally to angular velocity integration. The required scaling is $W^{\text{in}} \propto \cot\delta$, which means clones with larger phase shifts need weaker velocity input.

From the previous subsections, it is clear that if we have clones with phase-shifted outputs, and if their firing rates are differentially modulated, we end up with a term that makes the bump move. The additive velocity input is what achieves this differential modulation. Then, due to the ReLU activation, this results in a multiplicative effect when the differential input is small (see subsection 2.5).

The key constraint is that each clone pair's contribution to the bump velocity must be the same, regardless of its phase shift $\delta$. From equation (55), a clone pair with phase shift $\delta$ and differential gain $m$ contributes $C_{\text{eff}} = mJ_E \tan\delta$ to the odd component. If we want $C_{\text{eff}}$ to be the same for all pairs, then the velocity-driven gain imbalance $m$ must scale as $m \propto \cot\delta$. Since $m$ is proportional to the velocity input weight, this gives the velocity input scaling rule $W^{\text{in}} \propto \cot\delta$. Clones with larger phase shifts have larger recurrent weights (scaled by $1/(2\cos\delta)$) but need proportionally weaker velocity inputs. This is the origin of the $\cot(\Delta)$ scaling rule described in the main text.

##### 4.4.4 Balanced noise

The summed-output rule constrains only the *sum* of clone weights onto each postsynaptic neuron. This means that individual clone weights can vary – even substantially – so long as the noise sums to zero across the clone group. This is the mechanism by which hidden symmetries tolerate single-neuron heterogeneity.

The clone construction also admits balanced noise. Suppose

$$\widetilde{W}_{i,j\mu} = \overline{\widetilde{W}}_{i,j\mu} + \eta_{i,j\mu}, \qquad \sum_{\mu\in\mathcal{C}(j)} \eta_{i,j\mu} = 0. \tag{58}$$

Then the summed output from the clone group onto postsynaptic unit $i$ is unchanged. Single-clone weights can therefore look highly heterogeneous even though their group sum is exact.

If the clone group is further subdivided into phase-matched subsets (main text Figure 5), then the noise must balance within the relevant subset to preserve that subset's net phase shift. This is the precise sense in which hidden symmetries can be invisible at the single-neuron level but reappear after averaging across the correct clone group. Empirically, we find that the amplitude of the balanced weight noise can be about the same order as the amplitude of the structure weight component before affecting off-manifold stability.

### 5 Connection to canonical HD models

This section shows that the classical 2-ring and 3-ring HD models from the theoretical neuroscience literature are special cases of the dynamical clone framework. This unifying perspective reveals that these seemingly distinct architectures all implement the same underlying mechanism – counterbalanced phase shifts with appropriate scaling – and differ only in the number and arrangement of their clone populations.

Within theoretical neuroscience, the canonical ring attractor model reviewed in Section 2 has driven much of the conceptual understanding of head direction systems. Biologically, however, having a velocity-dependent term directly in the connectivity (the $\sin$ term in the kernel) requires synaptic weights that change in real time [14], which is difficult to implement in neural hardware. To circumvent this, two popular architectures – which we call the 2-ring [23] and 3-ring models [16] – use separate populations of neurons that receive angular velocity input and convert it into an effective asymmetric connectivity via their phase-shifted projections.

From the perspective of the clone construction developed above, these classical architectures are special cases of the more general framework:

**2-ring model [23]:** Starting from an $N$-neuron ring, clone every neuron once to get $2N$ neurons, and assign all clones identical phase shifts of $\pm\delta$. Collect all neurons with phase shift $+\delta$ and call them the "left ring", and collect all neurons with phase shift $-\delta$ and call them the "right ring." This is exactly the counterbalanced clone-pair construction of the previous section, applied uniformly to every neuron. When the left ring receives a stronger velocity input than the right ring (or vice versa), the differential gain produces the odd connectivity component that moves the bump.

**3-ring model [16]:** Same as above, but go from $N$ to $3N$ neurons with phase shifts $+\delta$, $-\delta$, and $0$ for each clone triplet. The zero-phase-shift clones form the "center ring," which maintains the bump in the absence of velocity input. The $\pm\delta$ clones form the left and right rings, which are activated during turns to drive the bump. This architecture provides additional flexibility because the center ring can be tuned independently from the side rings.

In both cases, the phase shifts, scaling rules, and velocity-input requirements follow directly from the general clone framework. The key insight is that the 2-ring and 3-ring models assign the same phase shift to every neuron within a ring, whereas the dynamical clone construction allows each clone group to have its own phase shift. This additional flexibility is what enables the heterogeneous connectivity observed in the fly connectome while preserving continuous attractor dynamics.

### A Reduction of firing rate equation to low modes

Differentiating the ansatz (4), we get

$$\tau\dot{h} = \tau\dot{h}_0 + \tau\dot{\rho}\cos(\theta-\psi) + \tau\rho\sin(\theta-\psi)\dot{\psi}, \tag{59}$$

$$= \tau\dot{h}_0 + \cos\theta\left[\tau\dot{\rho}\cos\psi - \tau\rho\dot{\psi}\sin\psi\right] + \sin\theta\left[\tau\dot{\rho}\sin\psi + \tau\rho\dot{\psi}\cos\psi\right]. \tag{60}$$

Similarly, for the choice of input (2), which is peaked at $\tilde{\psi}$, we have

$$I(\theta-\tilde{\psi}) = I_0 + \cos\theta(I_1\cos\tilde{\psi}) + \sin\theta(I_1\sin\tilde{\psi}). \tag{61}$$

And for the terms containing the convolution with $W(\theta-\theta')$, we have

$$\int d\theta'\, W(\theta-\theta')[h(\theta')]_+ = W_0 g_0 + \cos\theta\,[W_1 g_c - C g_s] \tag{62}$$

$$+ \sin\theta\,[W_1 g_s + C g_c]\,, \tag{63}$$

where $g_i$ are defined as the mode projections of the nonlinearity as follows:

$$g_0(h_0,\rho,\psi) = \int \frac{d\theta'}{2\pi}\; [h(\theta',t)]_+\,, \tag{64}$$

$$g_c(h_0,\rho,\psi) = \int \frac{d\theta'}{2\pi}\; \cos\theta'\;[h(\theta',t)]_+\,, \tag{65}$$

$$g_s(h_0,\rho,\psi) = \int \frac{d\theta'}{2\pi}\; \sin\theta'\;[h(\theta',t)]_+\,. \tag{66}$$

Equating for the different modes, we get,

$$\tau\dot{h}_0 = -h_0 + I_0 + W_0 g_0. \qquad \text{DC mode,} \tag{67}$$

$$\tau\dot{\rho}\cos\psi - \tau\rho\dot{\psi}\sin\psi = -\rho\cos\psi + I_1\cos\tilde{\psi} + [W_1 g_c - C g_s] \qquad \text{Cosine mode,} \tag{68}$$

$$\tau\dot{\rho}\sin\psi + \tau\rho\dot{\psi}\cos\psi = -\rho\sin\psi + I_1\sin\tilde{\psi} + [W_1 g_s + C g_c] \qquad \text{Sine mode.} \tag{69}$$

To arrive at the equations for the phase and amplitude, we have

$$\tau\dot{\rho} = \cos\psi \times \text{Eq.}(68) + \sin\psi \times \text{Eq.}(69), \tag{70}$$

$$\tau\rho\dot{\psi} = \cos\psi \times \text{Eq.}(69) - \sin\psi \times \text{Eq.}(68). \tag{71}$$

We therefore have,

$$\tau\dot{\rho} = -\rho + I_1\cos(\psi-\tilde{\psi}) + W_1(\cos(\psi)g_c + \sin(\psi)g_s) + C\,(\sin(\psi)g_c - \cos(\psi)g_s) \tag{72}$$

$$\tau\rho\dot{\psi} = I_1\sin(\tilde{\psi}-\psi) + W_1(\cos(\psi)g_s - \sin(\psi)g_c) + C(\cos(\psi)g_c + \sin(\psi)g_s). \tag{73}$$

For simplifying computations, we define

$$g_1(\rho, h_0, \psi) = e^{-i\psi}(g_c + ig_s), \tag{74}$$

whence we get the following equations

$$\boxed{\begin{aligned} \tau\dot{h}_0 &= -h_0 + I_0 + W_0 g_0 &(75)\\ \tau\dot{\rho} &= -\rho + I_1\cos(\psi - \tilde{\psi}) + W_1\mathfrak{Re}(g_1) - C\mathfrak{Im}(g_1) &(76)\\ \tau\rho\dot{\psi} &= I_1\sin(\tilde{\psi} - \psi) + W_1\mathfrak{Im}(g_1) + C\mathfrak{Re}(g_1), &(77)\end{aligned}}$$

which are (9)-(11).

### B Useful integrals for the bump regime

Here, we evaluate $g_0$ and $g_1$. For $0 < \theta_c < \pi$, define

$$\cos\theta_c = -\frac{h_0}{\rho}, \tag{78}$$

so that in the bump regime,

$$[h(\theta)]_+ = \rho\,[\cos(\theta - \psi) - \cos\theta_c]_+\,. \tag{79}$$

After shifting $\phi = \theta - \psi$, the active region is $|\phi| < \theta_c$. Therefore,

$$\begin{aligned} g_0 &= \rho\int_{-\pi}^{\pi}\frac{d\phi}{2\pi}\,[\cos\phi - \cos\theta_c]_+ &(80)\\ &= \rho\int_{-\theta_c}^{\theta_c}\frac{d\phi}{2\pi}\,(\cos\phi - \cos\theta_c) &(81)\\ &= \rho\,\frac{\sin\theta_c - \theta_c\cos\theta_c}{\pi}, &(82)\end{aligned}$$

which is (19). Likewise,

$$\begin{aligned} g_1 = e^{-i\psi}(g_c + ig_s) &= \rho\int_{-\pi}^{\pi}\frac{d\phi}{2\pi}\,e^{i\phi}\,[\cos\phi - \cos\theta_c]_+ &(83)\\ &= \rho\int_{-\theta_c}^{\theta_c}\frac{d\phi}{2\pi}\,\cos\phi\,(\cos\phi - \cos\theta_c) &(84)\\ &= \rho\,\frac{1}{2\pi}\left(\theta_c - \frac{\sin 2\theta_c}{2}\right), &(85)\end{aligned}$$

which is (20). The imaginary part vanishes because the integrand is odd.

### C Derivation of the $\theta_c$ equation

Differentiating $\cos\theta_c = -h_0/\rho$ gives

$$\sin\theta_c\,\dot{\theta}_c = \frac{\dot{h}_0 + \cos\theta_c\,\dot{\rho}}{\rho}. \tag{86}$$

Substituting (21) and (22) then yields (24).

### D Clone-pair identity and gain scaling

For completeness, the counterbalanced clone-pair identity used in the main text can be checked in one line:

$$\frac{A}{2\cos\delta}\cos(\Delta - \delta) + \frac{A}{2\cos\delta}\cos(\Delta + \delta) = A\cos\Delta. \tag{87}$$

If the two clones carry gains $1 \pm m$, then

$$\frac{A(1+m)}{2\cos\delta}\cos(\Delta-\delta)+\frac{A(1-m)}{2\cos\delta}\cos(\Delta+\delta) \tag{88}$$
$$= A\cos\Delta + mA\tan\delta\,\sin\Delta. \tag{89}$$

So the odd coefficient is $C_{\text{eff}} = mA\tan\delta$. Equivalently, if one writes the structured recurrent amplitude of a single shifted branch as $w_r = A/(2\cos\delta)$, then

$$w_r\sin\delta = \frac{A}{2}\tan\delta, \qquad C_{\text{eff}} = 2m\,w_r\sin\delta. \tag{90}$$

This is the direct dictionary between the continuum expression in this note and the finite-network main-text rule $g = w_r\sin(\Delta)$, with main-text $\Delta$ equal to the present phase shift $\delta$. If $m = \kappa(\delta)\omega$, then the reduced phase equation implies

$$\tau\dot{\psi} = \kappa(\delta)A\tan\delta\, f_1(\theta_c)\,\omega. \tag{91}$$

Therefore, maintaining a fixed integration gain requires $\kappa(\delta) \propto \cot\delta$ up to a common prefactor.

### E Remark on exact versus approximate cloning and slope rescaling

Equation (33) is exact only for constant positive rescalings. If $v = v(t)$, then

$$\tau\frac{\partial(vh)}{\partial t} = v\,\tau\frac{\partial h}{\partial t} + \tau\dot{v}\,h, \tag{92}$$

so a time-dependent gain produces an extra drive proportional to $\dot{v}$. This is why the dynamic-clone picture is exact for constant gains and approximate for slowly varying gains.

Likewise, the summed-output clone equivalence in Section 4.1 is exact when the clone constraints (46)–(47) hold and the clones within a group are initialized to share the same firing rate as their parent unit (which they will then maintain in the absence of input). In biological circuits and trained RNNs, one should expect these conditions to hold only approximately. The value of the exact construction is therefore not that biology must literally instantiate it, but that it shows the heterogeneities highlighted in the main text are consistent with exact continuous-attractor implementations.

Finally, some predicted amplitude scalings need not appear directly as anatomical weight scalings. For a threshold-linear unit with activation function $\phi(x) = s\,[x]_+$, multiplying all outgoing weights from that unit by a factor $\lambda$ is functionally equivalent to multiplying its slope $s$ by $\lambda$. Thus part of the required rescaling can be hidden in effective single-unit gain rather than in synapse counts alone.
